# The dynamic response of the bacterial flagellar motor to its direct intracellular input signal

**DOI:** 10.1101/2025.10.28.684865

**Authors:** Alina M. Vrabioiu, Basarab G. Hosu, Aravinthan D. T. Samuel

## Abstract

The bacterial flagellar motor drives bacterial swimming and chemotaxis by rotating helical flagellar filaments. When *Escherichia coli* navigates chemical gradients, the motor switches from counterclockwise (CCW) during forward swimming to clockwise (CW) during direction-changing tumbles. The motor responds indirectly to extracellular chemosensory input to membrane-bound chemoreceptors using an intervening intracellular signaling pathway. How the motor responds to its direct input signal – the diffusible messenger CheY-P – remains poorly understood. Steady-state motor measurements have been modeled as an allosteric switch between CCW/CW states that depend on mean CheY-P levels. Allosteric models have suggested that as many as 20 CheY-P molecules can be bound to the motor when it switches rotational direction. But steady-state models cannot predict the sensitivity of the motor to dynamic changes in CheY-P that essentially modulate chemotactic behavior. We present an optogenetic reagent that precisely controls the direct dynamical input signal to the motor. We designed a “caged” molecule, Opto-CheY, that is transiently activated by photon absorption. We find that activation and binding of 1-3 CheY-P molecules is sufficient to switch the motor from the CCW to CW state. The sensitivity of the motor to small changes in CheY-P occupancy helps resolve a long-standing paradox about the high sensitivity of the chemotactic response to external sensory input. Optogenetic biochemistry by light-activated uncaging of signal molecules is a new strategy to dissect information-processing in the living cell.

**Significance Statement:** Motile bacteria swim to better environments by modulating the rotation of the bacterial flagellar motor. How this motor responds to intracellular signaling activity is poorly understood. The physiologically-relevant response of the motor is to transient activation of intracellular signaling molecules on the sub-second time scale of bacterial decision-making. Here, we report the first optogenetic probe that targets *in vivo* the output module of the the chemotactic network. We demonstrate that the motor has high dynamical sensitivity to the binding of single intracellular signaling molecules. This solves a long-standing problem of high sensitivity and signal amplification in the bacterial chemotactic response.

---

**C**hemotactic behavior in *E. coli* is exquisitely sensitive. The binding of single chemoattractant molecules to membranebound chemoreceptors will modulate bacterial flagellar motor (BFM) rotation and swimming behavior. (1–4) The complete transformation from sensory input to motor response is a convolution of multiple biochemical steps from chemoreceptor activation to motor switching between counterclockwise (CCW) and clockwise (CW) rotation. The physics of chemoreception allows membrane-bound receptors to reliably count single ambient chemoattractant molecules. (1–3) However, it has been challenging to isolate and study specific information-processing steps and molecular interactions inside the cell. The last step — how the motor responds to its direct input signal, phosphorylated CheY (CheY-P) — remains poorly understood.

CheY-P is produced by CheA (kinase associated with chemoreceptors, Fig. 1a). CheY-P diffuses in the cytoplasm and binds to the C-ring, an intracellular component of the motor. (5, 6) The C-ring is an oligomer made of *M* = 34 − 44 identical subunits, each a heteromer assembled from FliM, FliN, and FliG proteins. (7–9) Thus, each CheY-P molecules diffuses and binds to one of *M* binding sites provided by FliM/FliN. (8, 10) The chemotactic response depends on a large conformational change by the C-ring when enough binding sites are occupied by CheY-P. (11, 12) The C-ring has different conformational states for CCW and CW motor rotation. (13, 14)

**Fig. 1.**
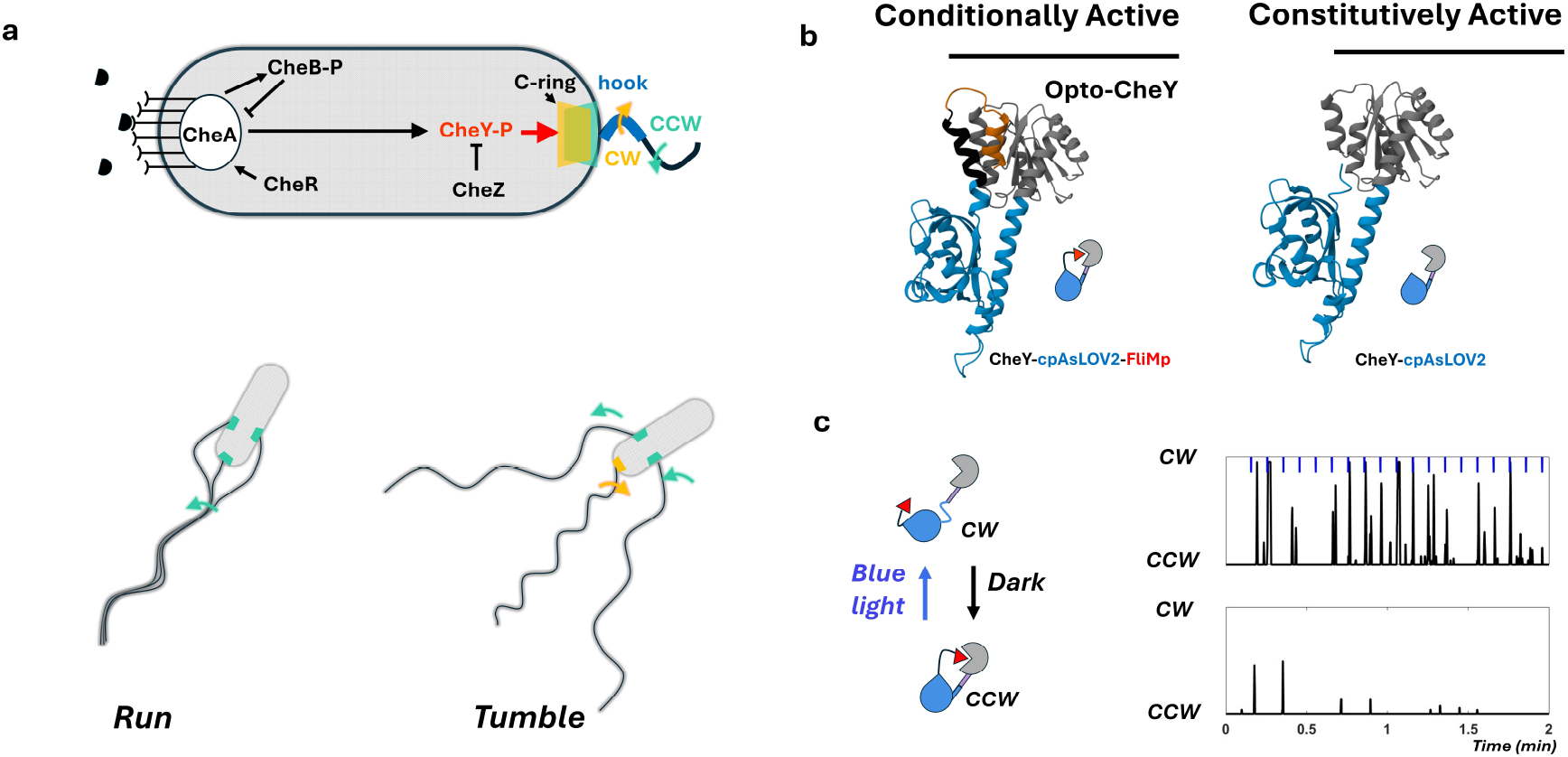
Opto-CheY, a ‘caged’, light-activated CheY. **a**. Diagram of key molecules in the chemotaxis network. Extracellular ligands (black semicircles) bind to receptors and modulate the activity of CheA, the kinase that phosphorylates the diffusible messenger CheY. CheA activity adaptation is mediated by the methylase CheR and de-methylase CheB. CheY-P binds to the C-ring and triggers a conformational change that results in BFM changing direction (CCW → CW). The swimming bacterium switches from a smooth run to a tumble. **b**. AlphaFold structure predictions of chimeric molecules. Grey is *E. coli* CheY, blue is cpAsLOV2, black is a short linker, and orange is the FliM peptide. Cartoons depicting the “caged” dark-state of Opto-CheY and the constitutively active control are shown with the same color code. **c**. Blue-light absorption triggers unfolding of the J*α* helix, and CheY activation by removal of the FliM plug from the CheY active site. Active CheY binds to the C-ring and promotes the rotational switch of the BFM from CCW → CW. Light pulses (blue dots, ~100 *µ* J/cm^2^) are applied every 6 seconds while monitoring BFM rotational state. Each response is a CW rotation (vertical black line) following a blue light pulse (blue dot) in an otherwise mostly CCW motor (bottom trace)

Cluzel et al.(15) measured the steady-state CW bias (fraction of the total time the motor spends in the CW state) for cells with different steady-state [CheY-P]. Higher CheY-P levels correlated with higher CW bias, following a steep concentration dependence (Hill coefficient ~10-20). (15, 16) The steep Hill curve is often interpreted as a mechanism for signal amplification. Near inflection, a small difference in steady-state CheY-P levels corresponds to a large difference in steady-state CW bias.

However, steady-state measurements inform the thermodynamics of motor switching at equilibrium, not how the system responds to the transient perturbations of sensory transduction that are essential for chemotaxis up chemical gradients. To probe the kinetic structure of chemotactic signal processing, one needs to probe the motor response to defined perturbations of sensory input or intracellular signaling.

How sensitive is the motor to its direct input signal? How many CheY-P actively bind to the motor when it switches from CCW to CW? The smallest input signal that evokes a CCW → CW switch should occur in a motor that is “dark-adapted” to low CheY-P levels. Analogous questions about the smallest quantal stimulus that triggers a behavioral response have been asked in visual neuroscience. (17) How many photons are absorbed by the dark-adapted retina to see the dimmest flash? (18–20) How many photons are absorbed by the dark-adapted rod cell to evoke an electrical response? (21) Visual neuroscientists decisively answered these questions to establish single-photon sensitivity in vertebrate vision.

To date, it has only been possible to indirectly change CheY-P levels using external chemosensory input. But any effect of external input on internal CheY-P is confounded by the intracellular signaling network. Cell-to-cell variability and stochasticity throughout this network adds further uncertainty when interpreting the motor response to external input. (22, 23) Here, we present an optogenetic approach to directly probe the sensitivity limits of the bacterial flagellar motor to its input signal.

## Results

Optogenetic tools allow light-activated perturbations of cellular signals. (24) We sought an optogenetic approach to directly manipulate CheY-P in the living cell. We wanted to decouple CheY-P activity from the endogenous signaling network, activate CheY-P with controlled light pulses, and monitor the bacterial flagellar motor (BFM) response in real-time.

### Designing Opto-CheY

Chemotactic signaling processing in *E. coli* works on the ~1-3 sec time-scale, the range of run lengths in free-swimming cells. (25–27) For bacterial chemotaxis, a suitable optogenetic probe must exhibit a light-activated conformational change that recovers its dark-state within this time-scale. (28, 29) We selected the LOV2 domain from *Avena sativa* (AsLOV2). The AsLOV2 domain contains an *α*-helix (J*α*) that unfolds upon absorbing a blue photon (~450 nm). (30) Wild-type AsLOV2 recovers its dark state with ~50 sec half-life. (30) Rapidly-refolding variants have been identified, including AsLOV2_N449S_ (<1 sec recovery (31)) and AsLOV2_V416T_ (~2 sec recovery (32, 33)).

Our goal was to design “caged” CheY by chimeric fusion with AsLOV2 (Fig. 1b). We wanted a molecule that would uncage when the J*α* helix unfolds upon photon absorption. To build a stable chimeric structure, protein orientation can be constrained by fusing adjacent domains at junctures within the same *α*-helix. (34) By adjusting the length of the *α*-helical juncture, the relative position and orientation of adjacent domains can also be fine-tuned. Here, we fused the J*α* helix of circularly permuted AsLOV2 (cpAsLOV2) (35, 36) to the C-terminal *α*-helix of CheY.

We sought a dark-state structure where a low-affinity inhibitory peptide would effectively cage CheY by plugging its motor binding site. For the inhibitory peptide, we chose 16 amino-acids from FliM, the endogenous binding partner of CheY, which we fused to the C-terminus of cpAsLOV2. (10) We hypothesized that photon-triggered unfolding of the J*α* helix would release the constraint on CheY and AsLOV2 orientation — this would remove the inhibitory peptide and expose the CheY active site. We used AlphaFold (37) to test the length of the *α*-helical juncture until we found a prediction where the CheY active site is plugged in the dark-state and unplugged in the lit-state. We call this molecule Opto-CheY (Fig. 1b).

### Testing Opto-CheY in an optimal bacterial strain

CheY does not bind the C-ring to promote CW rotation unless it is activated by phosphorylation to CheY-P. CheA is the kinase that acts on CheY. CheB is the de-methylase that reduces CheA activity. CheZ is the phosphatase that dephosphorylates CheY-P (Fig. 1a). (38, 39) In cells lacking CheZ and CheB, intracellular CheY is completely phosphorylated. (40) Thus, to maximize motor sensitivity to optogenetic activation, we expressed Opto-CheY in a Δ*cheB*Δ*cheZ*Δ*cheY* strain background.

Previous steady-state measurements found that the motor is most sensitive to CheY-P in its normal endogenous range, 2-4 *µ*M. (15) We titrated Opto-CheY levels using insulated, variable-strength bacterial promoters (41) and used fluorescence correlation spectroscopy (FCS) to verify that CheY expression is within endogenous range (Fig. S1).

Using targeted mutations of promoter and/or ribosome binding sites, we fine-tuned Opto-CheY expression to an optimal level where optogenetic stimulation spans the full dynamic range of motor response, from nearly fully CCW rotation to fully CW rotation (Tables S1, S2).

To determine the rotational direction of single motors we used the tethered cell approach. (42) To tether cells directly by hooks (Fig. 1a) we developed a “sticky” hook approach. Specifically, we increased the number of positive charges in a surface exposed loop of the flagellar hook protein FlgE. This promotes the hook’s electrostatic interaction with the negatively charged glass surface (Methods). (43, 44) We tethered cells that express Opto-CheY_N449S_ (strain number 638, Table S1) onto microscope coverslips. Pulses of blue light (~100 *µ* J/cm^2^) trigger single CCW → CW switching events in motors that rotate exclusively or almost exclusively in the CCW direction (Fig. 1c, Fig. S2). In contrast, cells that do not express Opto-CheY (strain number 657, Table S1) never respond with CW rotation (Fig. S2, Movies 1-6). These observations establish that blue light activates Opto-CheY to bind to the motor and promote CW rotation.

Tethered cells that express Opto-CheY_N449S_ have motors that are nearly exclusively CCW in dark-state (average CW bias 0.011 *±* 0.0032 for strain number 638, Table S1). We attribute low but non-zero CW bias in dark-state to dark-state equilibrium between folded and unfolded AsLOV2. (45, 46) This results in a population of Opto-CheY molecules that are active in the absence of optogenetic stimulation. If the motor reversals were indeed due to dark-state equilibrium, decreasing Opto-CheY expression should decrease the absolute amount of Opto-CheY molecules active in the dark and decrease the initial CW bias. When Opto-CheY expression level is decreased by ~20% via a base pair change in the ribosome binding site (47) dark-state CW bias decreased to 0.00185 *±* 0.00064 (strain number 630, Fig. S3). Conversely, when Opto-CheY expression is increased by ~33% with changes in both the ribosome binding and promoter sites dark-state CW bias increased to 0.085 *±* 0.022 (strain number 670, Fig. S3). The CW bias of cells that express CheY and cpAsLOV2 without the inhibitory peptide (strain number 658, Table S1) is 0.982 *±* 0.0035 (Fig. 1b, Fig. S3). We conclude that the entire range of the motor’s response — from fully CCW in cells with caged Opto-CheY in the dark-state to fully CW in cells with constitutively-active CheY-cpAsLOV2 — can be spanned by optogenetic control.

### Quantal stimulus at response threshold

The motor switches to CW rotation when enough CheY-P are added to the C-ring. (48) Opto-CheY is transiently activated when it absorbs a single blue photon. By measuring when the motor ‘sees’ an optogenetic pulse by switching from CCW → CW, we can estimate the minimum number of activated Opto-CheY that are added to the motor to trigger a switch.

In visual neuroscience, Hecht, Shlaer, and Pirenne (18) measured the minimum quantal stimulus at the retina – the number of rhodopsin molecules that must be activated by photons – for a dark-adapted human to see a dim flash of light. To do this, they made ingenious use of Poisson statistics. When the dark-adapted eye is exposed to dim flashes of light, each rhodopsin within the field of illumination has a small probability (*p* ≪ 1) of absorbing a photon and contributing to ‘seeing’. Because there are many rhodopsins (*N* ≫ 1) that can potentially be activated, the probability that a specific number of rhodopsins are activated must follow Poisson statistics. A given light flash delivers a total number of photons per unit area (*F*, photons/cm^2^) and activates on average a certain number of rhodopsins (*a* = *p · N* = (*const* · *F* · *σ* · Θ) ·*N*), *σ* and Θ are the absorption cross-section and quantum yield of rhodopsin, *const* accounts for both efficiency in signal transductions and photon dissipation. From this, Poisson statistics defines the probability that any specific number of rhodopsins (*k*) are activated:

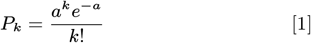

The probability that a flash evokes a behavioral response is the cumulative probability that the flash activates any number of rhodopsins above a threshold, *θ*:

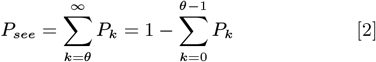

The average number of activated rhodopsins is proportional to flash intensity (*I* = *F/τ*, where *τ* is the duration of the light pulse). Rearranging, *a* = (*const* · *N* · *τ* · *σ* · Θ) · *I* = *α · I*, where *α* is constant for a given experimental system and

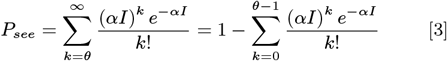

When the probability of seeing is plotted against the natural logarithm of flash intensity (ln *I*) a sigmoidal response curve will result. The slope of this curve at inflection depends strictly on *θ*. (49) The steeper the curve, the higher the quantal threshold needed to ‘see’ the flash.

Our experimental setup allows a similar approach. (18– 20, 50) Because the motor is nearly always in the CCW state in the dark, it tells us when it ‘sees’ a pulse with a CCW → CW switch. Each cell has ~1000-2000 Opto-CheY (Fig. S1) that are potentially activated by an optogenetic pulse (*N* ≫ 1). Each Opto-CheY has a small probability of being activated and diffusing and binding to the motor (*p* ≃ 0.005 ≪ 1, SI text). When a threshold number of Opto-CheY are added to the motor, they trigger a CCW → CW switch. By measuring motor response probability as a function of pulse intensity, we can leverage Poisson statistics to use Eq. 3 and estimate *θ* — the threshold, in our case the number of Opto-CheY molecules that must be added to the motor to trigger a response.

We tethered cells expressing Opto-CheY_V416T_ in the Δ*cheB*Δ*cheZ*Δ*cheY* background (strain number 636, Table S1). Here we used the Opto-CheY variant with ‘lit’ state half-life ~ 2 sec to lengthen the time Opto-CheY is in the active, signaling state. This strain has the lowest expression level that evokes motor responses (average initial CW bias is 0.0012 *±* 0.0004). We selected motors with initial dark-state CW bias close to 0.001 (equivalent 2-5 reversals per minute). Thus, the dark-state level of switching behavior in these motors, attributed to dark-state equilibrium between folded and unfolded Opto-CheY, is situated near the base of the response curve of previous steady-state experiments (15).

We applied blue-light flashes (2 ms) with geometrically increasing intensity (*I* = *I*_*min*_ *x, x* = {1, 2, 4, 8, 16, 32}, *I*_*min*_ flash delivers ~ 7 · 10^12^ blue photons/cm^2^, Fig. S4, Movies 7, 8). The frequency of the flashes is constant for a given intensity (0.25 Hz, every 4 sec, for *I*_*min*_). The interval between flashes was chosen such that each motor response is independent of the previous response (according to criteria outlined in SI text, Figs. S5, S6). The minimum interval that satisfies the criteria is a function of flash intensity; the stronger flash requires a longer time interval to recover the baseline (0.0667 Hz, every 15 sec for the maximum flash intensity, Fig. S4). At each intensity, we estimate response probability (*P* = # of flashes that trigger a CCW → CW switch divided by # total flashes). We plotted response probability versus the logarithm of relative flash intensity (Fig. 2).

**Fig. 2.**
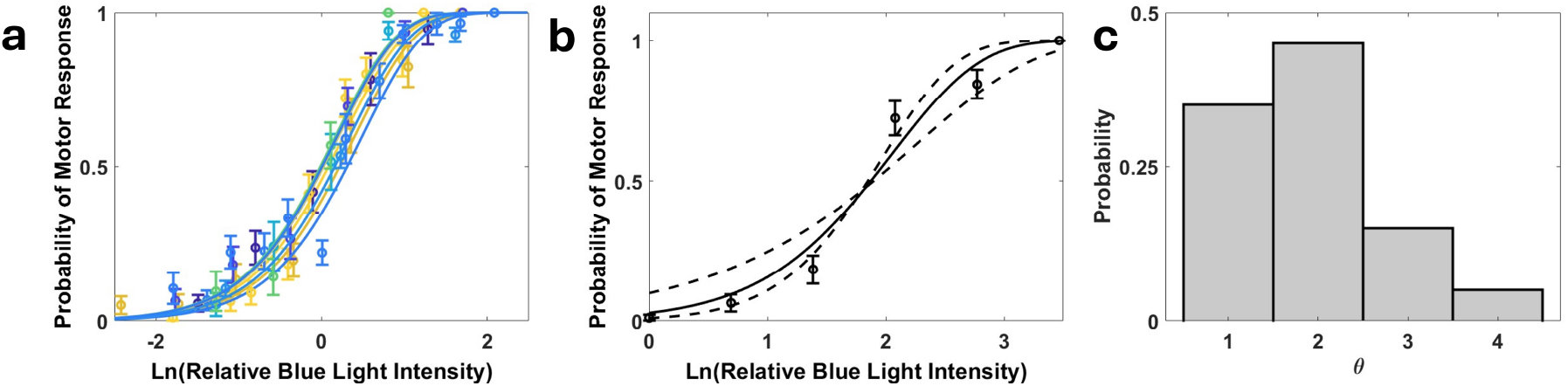
Probability of motor response. **a**. Opto-CheY_V416T_-driven motor responses. Each motor is flashed with a series of pulses of increasing relative intensity (*I* = *I*_*min*_ · *x, x* = {1, 2, 4, 8, 16, 32}, *I*_*min*_ delivers ~ 7 · 10^12^ blue photons/cm^2^). Response probability for each BFM at a given intensity is calculated as the number of responses divided by the number of flashes. P_see_ with *θ* = 2 is used to fit each dose response curve (equation 3). Ten individual BFM response curves and fits are shown translated horizontally relative to half-maximum response intensity. **b**. The dose response of one Opto-CheY_V416T_ motor is shown along with best fit curves for fixed values of *θ*: *θ* = 2 (black line), *θ* = 1 (shallower dashed black line), *θ* = 3 (steeper dashed black line). **c**. A histogram of most probable *θ* for 20 individual BFM response curves, corresponding to the fit with highest *R*^2^.

We fitted the data for each motor to Eq. 3 (SI text). Best fits are in the range *θ*=1-3 for all but one motor (Fig. 2c). Fig. 2a shows 10 individual motor responses (motors 1-10 in Table S5) along with corresponding *θ* = 2 fits. Individual responses/fits are translated horizontally so they align to the half maximal response and highlight the steepness of the curves (raw responses are shown in Fig. S8).

For single motors, the cumulative Poisson function is often fit to slightly different values of *θ* with similar *R*^2^ values (Table S5). For example, *θ*=2 is the best fit for the motor response shown in Fig. 2b, but *θ*=3 fit is a close second, suggesting measurement uncertainty of *θ ±* 1. In our model, *θ* is an estimate of the number of Opto-CheY that must be activated and added to the motor to reach the threshold of CCW → CW switching. Differences in *θ* can also be attributed to differences in dark-state Opto-CheY occupancy. Indeed, we found that motors that start at higher initial biases (motors 11-20 in Table S5) have higher response probability at the lowest blue light intensity (y-axis intercepts above 0.1 in Fig. S8, V416T panel) as well as flatter overall responses that are best fit with a *θ* = 1 (Table S5). Other Opto-CheY strains with higher dark-state CW bias were also best fit by shallowed cumulative Poisson functions with low *θ* values (strain number 638, Fig. S9c and its caption).

Opto-CheY_V416T_ recovers its dark-state in ~2 sec after activation. (32) Opto-CheY_N449S_ recovers faster, in <1 sec. (31) We measured similar dose-response curves using Opto-CheY_N449S_ (strain numbers 630 and 638, Table S1). Increasing light intensity led to increasing response probability with similar steepness (*θ*~2) (Figs. S8, S9). However, response probability with the shorter-lived probe did not always saturate close to 1 even with the strongest light pulses (Fig. S7). Rapid inactivation puts an upper bound on response probabilities. If Opto-CheY molecules refold before enough have time to bind the C-ring, no response can be evoked. Consistent with this idea, the motor can respond to every stimulation of Opto-CheY_N449S_ when longer light pulses of lower intensity are used, which effectively increases the life-time of the probe (Fig. S10).

### The Impulse Response of the Motor

In classic impulse response measurements, tethered cells were exposed to short pulses of chemoattractant from an iontophoretic pipette. (25, 26) These cells expressed wild-type CheY at endogenous levels and had higher initial CW bias. Following each iontophoretic pulse, CCW likelihood rises for ~1 sec, returns to baseline, falls below baseline, and finally returns to baseline after ~3 sec. This biphasic response reveals the time-scale of chemotactic signal processing. This response time-course is a mathematical convolution of all signaling events from chemoreceptor activation to motor switching.

Optogenetic activation of Opto-CheY allows us to measure the kinetics of the last event in the signaling pathway, the interaction of CheY-P with the C-ring. Because nearly every CCW → CW switch is light-evoked, each response is characterized by two time intervals. Latency is the interval between the optogenetic pulse and the CCW → CW switch (the time it takes for the flash to evoke a response). Duration is the interval between the first CCW → CW switch and the return to the CCW state (more precisely, the duration of the first CW interval following the flash, Fig. 3a). As in the classic impulse response experiments (25, 26), the average of single motor binary traces following blue light stimulation generates the probability of being in the CW state at a given time after the light pulse (Fig. 3a).

**Fig. 3.**
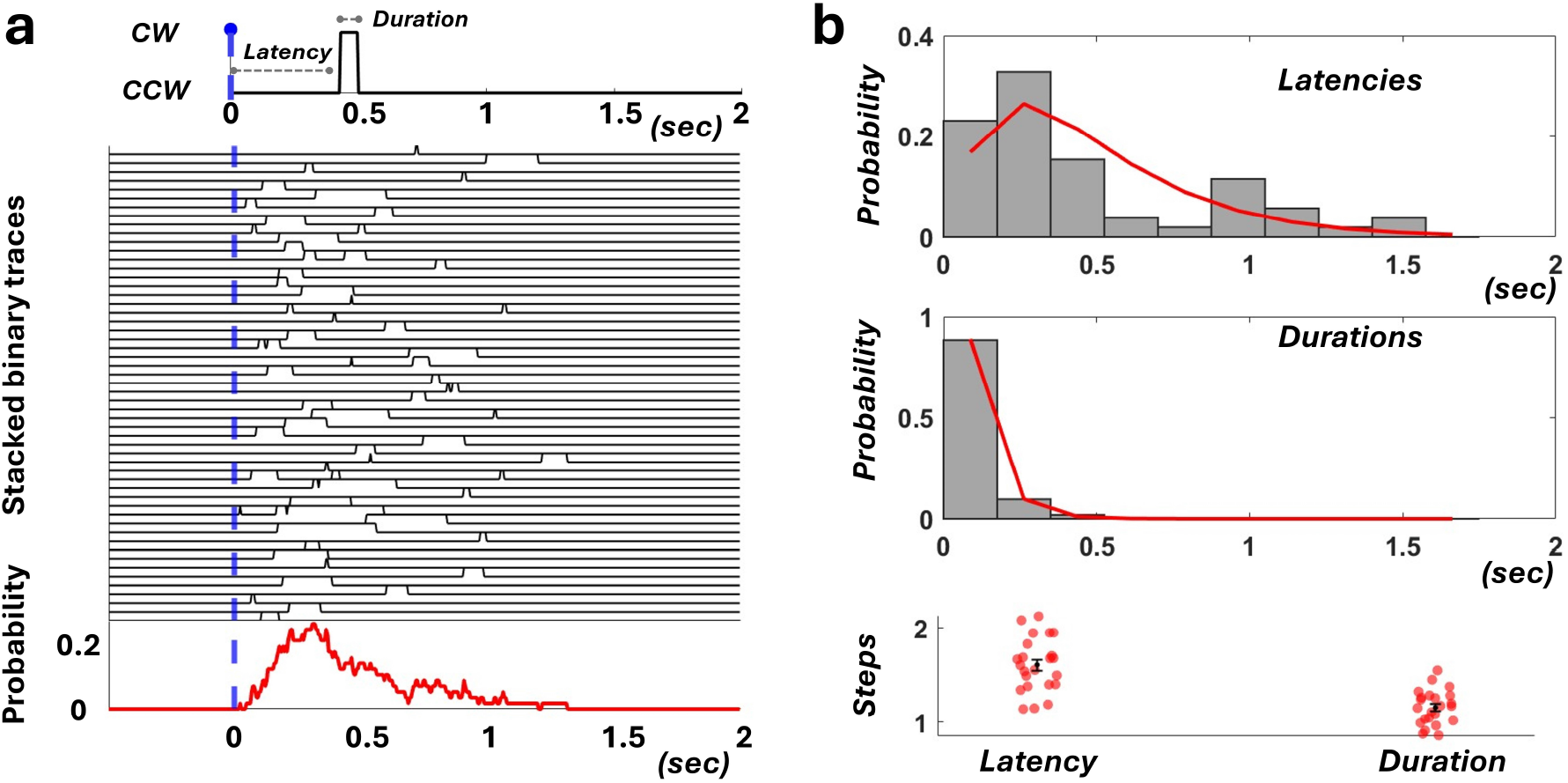
Impulse response of the motor. **a**. Each response is decomposed into the time it takes to respond (Latency), and the duration of the first CW interval (Duration). Opto-Che 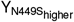 motor is subjected to 400 ~blue light pulses (*I* = *I*_*min*_ · *x, x* = 4) and responded 54 times (response probability=0.135). The 54 positive trials are aligned relative to the blue-light pulse, and averaged to generate the likelihood of being in the CW state at a given time after the pulse (bottom, red). **b**. Histograms of the 54 latencies and durations are shown (normalized to represent probabilities). Latencies are fit with a gamma function (shape parameter 2, *n* = 2). Durations are fit with an exponential function (shape parameter 1, *n* = 1). Different BFM responses are each fitted to estimate the number of rate limiting steps for latency and duration. Individual points, means, and standard errors are shown.

The statistics of the optogenetically-induced switching response reflect the stochastic nature of CheY-binding kinetics. Fluctuations in response latency (e.g., the time it takes to reach the CW state, latency in Fig. 3a) can be informative about elementary steps in the overall biochemical reaction. (51–53) An elementary, single-step reaction is a Poisson process with exponentially distributed completion times. For a multi-step reaction mechanism (multiple, sequential elementary steps) the distribution of overall reaction times is given by the convolution of its elementary steps duration distributions, dominated by the slowest steps. (53) For *n* comparable, rate-limiting steps, the probability distributions of total reaction times are best fit by gamma functions with shape parameter equal to *n*.

We collected statistics for latency and duration using cells that express Opto-CheY_N449Shigher_ (half-life <1 sec, strain number 638, Table S1) at different flash intensities. For the cell shown in Fig. 3a, 400 blue light pulses were applied (*I* = *I*_*min*_ · *x, x* = 4), and the motor responded 54 times (response probability = 0.135). Histograms of 54 response latencies and durations are shown in Fig. 3b. For a weak stimulus with low response probability (13.5% of flashes evoke a reversal), response latency is a significant proportion of the overall signal processing time (Fig. 3b).

Probability distributions for latency are poorly fit by single-exponential curves. Response latency is better fit by the gamma distribution. Assuming that *n* rate-limiting steps with similar kinetics occur in series, *n* can be estimated from the best-fit gamma distribution.

When gamma distributions are fit to response latency histograms, we estimate *n*~2 for most cells (Fig. 3b, bottom). One possibility is that response latency is the result of two rate limiting steps – such as the activation and successive binding of two Opto-CheY molecules to reach the CW state of the motor. However, the probability distributions for the first CW interval duration at low flash intensity are well-fit by single-exponential curves (Fig. 3b). Response duration might be the result of one rate-limiting step if only one Opto-CheY needs to unbind to return the motor to the CCW state.

Higher intensity flashes evoke higher motor response probabilities (Fig. 2, Figs. S8, S9) presumably because they activate a higher number of Opto-CheY molecules. Response latency and duration are also modulated by stimulus strength (Fig. 4a). Mean latency and duration for a stimulus with ~14% response probability (*x* = 4) are 0.5 *±* 0.05 sec and 0.11 *±* 0.013 sec, respectively. For a stimulus with ~68% response probability (*x* = 16), mean response latency drops to 0.18 *±* 0.015 sec and mean response duration rises to 0.16 *±* 0.016 sec (Fig. 4a). This trend is consistent across 24 motors that we studied with both weak and strong stimuli – the stronger the stimulus, the shorter the latency and the longer the duration of a response.

**Fig. 4.**
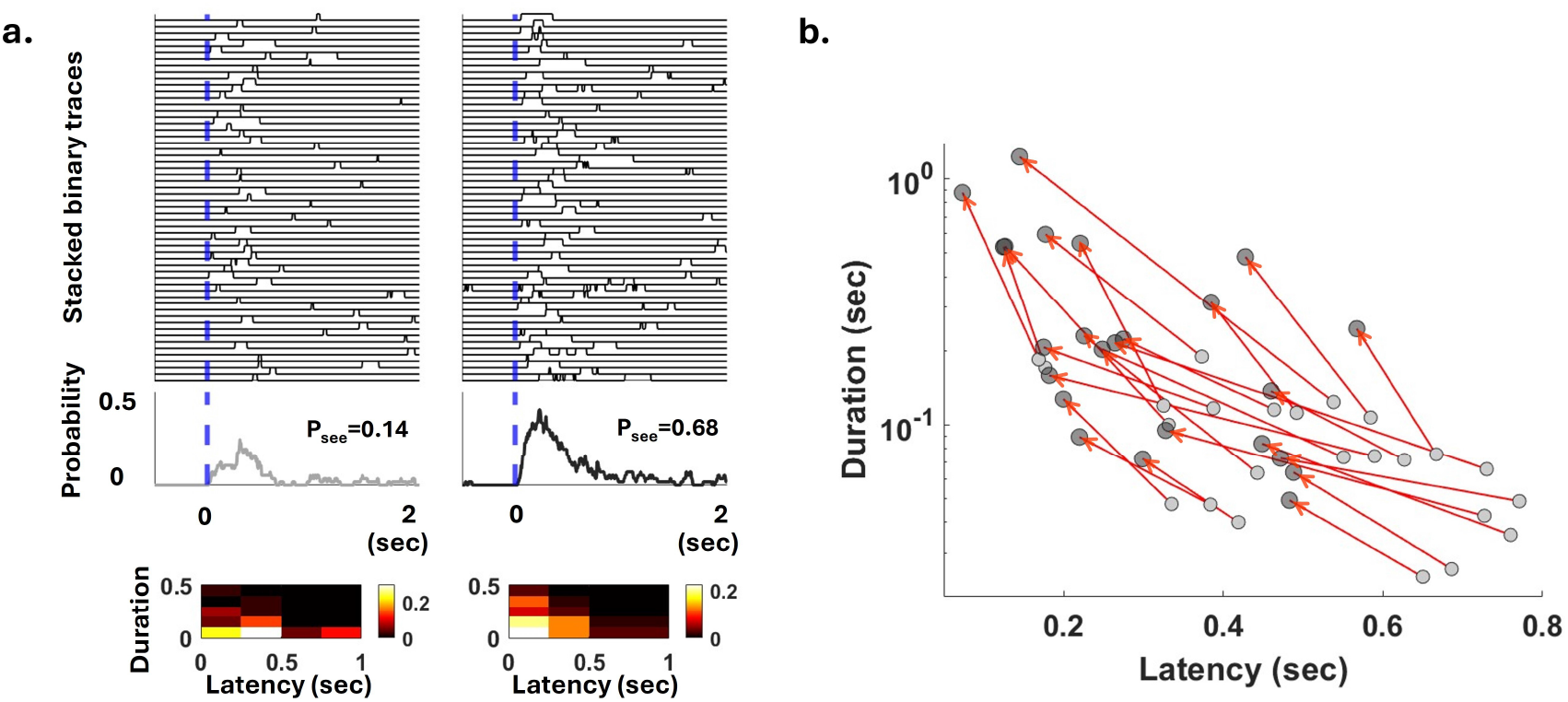
Motor response to weak and strong stimuli. **a**. Opto-Che 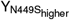A single motor is subjected to 400 weak light pulses (*I* = *I*_*min*_ · *x, x* = 4) and 120 strong pulses (*x* = 16). 57 responses for each stimulus are shown stacked, and aligned relative to the light pulse (blue line). Average motor responses for each stimulus is shown (gray for weak; black for strong). Below, the 57 latencies and durations are used to generate a joint probability distribution shown as a 2D-histogram, and color coded for probability as shown in the colorbar. **b**. Average latencies and durations for the weak (gray) and strong (black) stimulus are shown for 24 motors. Red line interconnect responses of each motor, pointing toward the strong stimulus.

Fig. 4b shows single motor responses at a given stimulus strength as points in the latency-duration space. Arrows connect the response to the weaker stimulus and the response to the stronger stimulus for single motors. Differences in the slopes of the lines reflect differences in individual motor dynamic responses. Increases in stimulus strength are correlated with decreases in latencies and increases in response duration, with different contributions across different motors. The same trend is observed in Opto-CheY_V416T_ motors (half-life ~2 secs, Fig. S11). However, the Opto-CheY_V416T_ motors take longer to return to the pre-stimulus baseline due to a longer lived perturbation.

## Discussion

The chemotactic signaling pathway in *E. coli* has long been noted for high gain and sensitivity – small changes in chemoreceptor occupancy cause large changes in bacterial flagellar motor (BFM) bias and swimming behavior. How the motor contributes to overall signal processing through its dynamical response has been poorly understood. Here, we studied the motor that is “dark”-adapted to low baseline CheY-P levels to probe the limit of molecular sensitivity. We report a dynamic sensitivity of the motor switching response to the binding of only 1-3 CheY-P molecules. In the regime of low baseline CheY-P levels, where motors rotate CCW, the high sensitivity of the motor to the binding of single signal molecules can contribute directly to signal amplification.

Previous theoretical and experimental work has suggested that the threshold of motor switching occurs near ~50% occupancy of CheY-P binding sites. (11, 12) However, the essential chemotactic response involves the adding and subtracting of small numbers of CheY-P molecules around the threshold of motor switching. Previous modeling has focused on the thermodynamics of conformational change and subunit coupling within the C-ring. (11, 54) The dynamics of CheY-P occupancy near the switching threshold is not well-understood. Diffusion-limited binding at endogenous [CheY-P] is fast (~ 10*/sec*). CheY-P also unbinds at a rate of ~ 10*/sec*. (11, 55, 56) A C-ring with *M* independent CheY-P binding sites can be modeled with *M* + 1 binding states (state *i* given by the number of occupied CheY-P binding sites). As the C-ring approaches the switching threshold from low occupancy, the kinetics will vary over time. As C-ring occupancy increases, the number of available binding sites decreases (site depletion), which should both increase the ‘on’ rate and reduce the ‘off’ rate. Thus, adding CheY-P molecules should become more rate-limiting as the motor advances from low occupancy to high occupancy states. Moreover, the C-ring is also thought to adaptively change the number of binding sites in response to sustained changes in CheY-P levels. (9) The motor is also thought to shed binding sites during CW rotation (as low as *M* = 34) and add binding sites during CCW rotation (as high as *M* =44).

The motor also adapts to changes in external load, as the mechanical load increases, more torque-generating stator units bind to the rotor (57), and CheY-P displays higher affinity for the C-ring. (58) Change in binding affinity can also affect the dynamical response. Changes in binding affinity might not affect the amount of binding occupancy near the switching threshold. However, an increase in binding affinity might affect how fast the switching threshold is reached after stimulation, reflected in response latencies.

In our experiments, the motor initially has very low CW bias and is presumably adapted to very low CheY-P levels. Tethered cells rotate slowly, in the limit of high mechanical load. Our estimates of *θ* in the cumulative Poisson functions (Fig. 2) and the non-exponential distribution of response latencies (Fig. 3) are both consistent with a small number of discrete rate-limiting steps to reach the switching threshold. We conjecture that motors in our experiments are adapted both to high mechanical load and dark-state conditions. The adapted motor has to bind enough unfolded Opto-CheY in the dark-state to be near the switching threshold, and thereby maximally-sensitive to optogenetically-induced increase in Opto-CheY levels.

In the regime of low baseline CheY-P levels, the high sensitivity of the motor to the binding of single signal molecules can contribute directly to signal amplification. Chemoreceptor-associated CheA molecules need only phosphorylate a small numbers of additional CheY molecules to trigger a tumbling response. The more CheY-P that is activated, the quicker the motor response. Two factors determine how many CheY-P must bind to the motor to evoke a CCW → CW switch: how many CheY-P are bound to the motor before the switch and how many more are needed to reach threshold. Response latency is similarly dependent on these factors. The stronger the stimulus, the shorter the response latency.

Optogenetic activation of CheY-P allowed us to study the dynamics of final reaction of the chemotaxis network. This approach can be extended to the swimming cell, offering a new way to test non-equilibrium theories of information-processing in bacterial chemotaxis. (59–62) An advantage of our pipeline is that it offers a flexible and modular way to design new optogenetic probes. We anticipate the extension of optogenetics to other proteins in the chemotactic signaling network (e.g., CheA, CheR, CheB). Our approach to inventing effective optogenetic probes is adaptable for *in vivo* dissection of intracellular signaling in diverse cell types.

## Supporting Information Appendix (SI)

### Materials and Methods

#### E. coli strains and plasmid construction

Motile MG1655 strain is the parent strain in all experiments. (63) Deletions of *cheB, cheZ, cheY, fliC, flgE*, and *fliK* were made sequentially, alone, and in combination (Δ*cheB*Δ*cheY* Δ*cheZ*) using the Datsenko and Wanner method with pKD3/pKD4 plasmids. (64) An 85 bp scar remained after Flp/FRT recombination to eliminate the antibiotic resistance genes. Deletions were verified using PCR and Sanger sequencing of the PCR products. Bacterial strains are shown in Table S1.

#### Opto-CheY construction

*cheY* DNA coding sequence was amplified from *E. coli* genomic DNA; cpAsLOV2 synthesized as a G block by Invitrogen (Table S4). Fragments with various linkers and truncations were assembled using the NEBuilder® HiFi assembly method (New England Biolabs) into the pEB2 plasmid backbone (Addgene #104007, kanamycin resistance and low-copy origin SC101 (65)) with a few modifications. Specifically, mutations were introduced to generate the series of pro1, pro3, proA, and pro5 promoters with increasing expression strength. (41) The ribosome binding sites used were a T7 RBS variant (GAAGGAGgT) which reduces protein expression by ~ 20% (47), and the wild type T7 RBS (Table S2). Mutations (N449S,V416T) were introduced via PCR (numbering as in AsLOV2 sequence, consistent with previous publications). Plasmids are shown in Table S2. Protein sequences are shown in Table S3.

#### Sticky hook construction

We based the design of “sticky hooks” on a previous insertion of the AviTag protein sequence into the flagellar hook protein FlgE (*FlgE*_*C*_*Avi*). (43, 44) There are 3 surface-exposed, negatively charged AviTag amino-acids (GLN**D**IF**E**AQKI**E**WHE, Fig. S12). We replaced these aminoacids with 3 positively charged ones (GLN**R**IF**R**AQKI**R**WHE) to promote interaction with the negatively charged glass surface (Table S3). The modified sequence was introduced via PCR into *flgE* amplified from genomic DNA to generate FlgE_AviRRR_ which was placed into the pTrc99A plasmid. (66)

#### Cell preparation and attachment to glass slides

Cells for microscopy were inoculated in tryptone broth (TB) with necessary antibiotics at 1:1000 dilution and grown at 27° C till OD_600_= 0.55 to 0.6. Leaky expression of FlgE_AviRRR_ in the pTrc99A plasmid was sufficient for attachment in the MG1655 strains. Cells were spun down, washed and re-suspended in motility buffer with no salt (10 mM K_2_HPO_4_/KH_2_PO_4_, 100 *µ*M EDTA, 10 mM lactate, pH=7). Cells were then added to channel slides (made by combining a glass slide, 2 layers of sticky tape, and coverslip) and allowed to settle for 5-10 min on the coverslip. After thorough washing (~ 0.5-1.0 mL of motility buffer), the tethered cells were used for experiments.

#### Microscope setup and imaging assay

We used a Nikon Eclipse Ti-U microscope and an Apo 60X oil objective (NA 1.49). The epifluorescence port was used to attach the blue-light LED for optogenetic stimulation. Neutral density filters (NDx reduce light intensity by 1*/x, x* = 1, 2, 4, 8, 16, 32) were used to modulate the intensity of the blue light pulse. Phase contrast images were acquired using a FLIR camera at 200 frames per second. A yellow filter was placed in front of the phase contrast lamp to filter out unwanted blue light during image acquisition. The LED-generated blue light pulse transiently increases the intensity of the frames it overlaps during phase contrast acquisition (1-2 frames, depending on pulse positioning and intensity). This pinpoints the localization of blue light pulses during image analysis. Trains of short pulses (2-5 msec) at each light intensity were applied every 2-15 sec.

#### Image processing and data fitting

Phase contrast movies were analyzed using ImageJ and Matlab with custom software. Each frame was assigned a rotational direction (CCW/CW), and CW bias was calculated as the number of CW frames divided by the total frames in a given time interval. Response probabilities were computed as *P* = # of flashes that trigger a CCW → CW switch divided by # total flashes of a given intensity.

## ACKNOWLEDGMENTS

This work was supported by NSF (award number 2146519) and Dean’s Competitive Fund at Harvard University. We are grateful to Thierry Emonet, Phillipe Cluzel, and members of the Samuel lab for discussions and support.

## Supporting Information

### Text Supplementary methods

#### Fluorescence correlation spectroscopy (FCS)

MG1655 Δ*cheB*Δ*cheY* Δ*cheZ*Δ*f lgE*Δ*fliC*Δ*f liK* cells express a CheY-pMag-mNeonGreen fusion using the proA promoter (strain number 649, Table S1). The plasmid backbone has kanamycin resistance and the low-copy origin SC101. (1) pMag is a variant of the fungal VVD photoresceptor of similar size and fold to AsLOV2 (2), and mNeonGreen was chosen for its increased photo-stability and fast maturation time.(1, 3) Cells were grown at 27° C till OD_600_=0.6, and incubated at 10° C to ensure full maturation of the reporting fluorescent protein. Cells were then washed in motility buffer and attached to the poly lysine treated coverslip of a 35 mm MatTek dish. Zeiss LSM 880 confocal (40X water objective, *NA* = 1.1) and the Zeiss FCS software were used to acquire data at the Harvard Center for Biological Imaging. Alexa488 at a concentration of 30 nM was used to calibrate the system. We analyzed only the cells with ≤15% fluorescent intensity decrease during the 2 sec acquisition time. We determined the average number of fluorescent molecules in the measured *E. coli* volume (Fig. S1). The width of *E. coli* is smaller than the axial dimension of the confocal volume (~ 844 nm with confocal pinhole at 0.66 airy units). To convert to concentration, we considered a lower (500 nm), and upper (750 nm) bound for the width of *E. coli* (Fig. S1). Bleaching of mNeonGreen results in a slight underestimate of the number of molecules in the measured volume and of the CheY concentration.

#### Blue light illumination

The blue light LED was mounted on a side port on the Nikon Ti-U microscope, with neutral density filters that could be slided in and out of the light path (NDx, reduced light intensity by 1*/x*). We measured the maximal LED power output used in our experiments to be ~ 45*µ*W. For a 2 msec blue light pulse, that comes to ~ 100*nJ* which corresponds to ~ 2.2 · 10^11^ blue light photons. The field of view (FOV) for our objective has a diameter of ~ 400*µ*m and an area of ~ 10^−3^*cm*^2^. This gives a maximal energy output of ~ 2.2 · 10^14^ photons*/cm*^2^; the minimal output is 1/32 of that which comes to 7 · 10^12^ photons*/cm*^2^. The probability of CheY activation at maximal energy is *p* = const · *σ* · Θ · *F*_*max*_. For Θ ~ 40%, *σ* ≃ 5.4 · 10^−17^ cm^2^, and *const* = 1, *p*_*max*_ ≃ 0.005 ≪ 1.

#### Data Fitting

Individual motor dose-response data was fitted to the cumulative Poisson probability functions (*P*_*see*_), plotted below for *θ* = 1 − 3, *a* is the average number of activated Opto-CheYs.

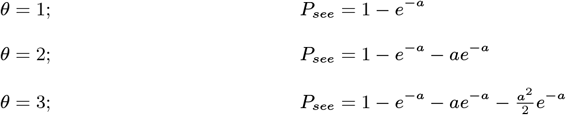

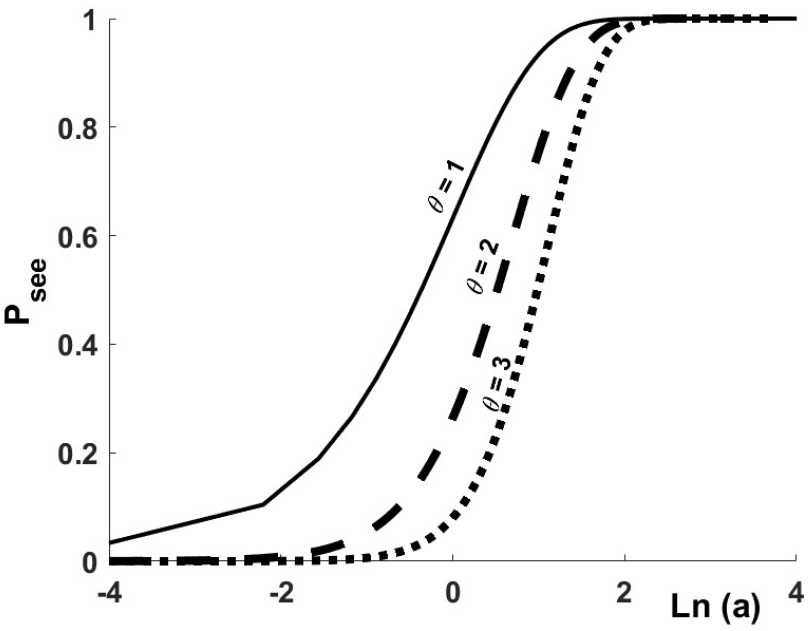

For our experiments *a* = *αI* = const · *N* · *σ* · Θ · *F*_*min*_ · *x*, 1 ≤ *x* ≤ 32. Since *F*_*min*_ ~ 7 · 10^12^ blue photons/cm^2^, *N* ~ 2000 Opto-CheY molecules, Θ ~ 40%, and 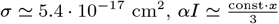.

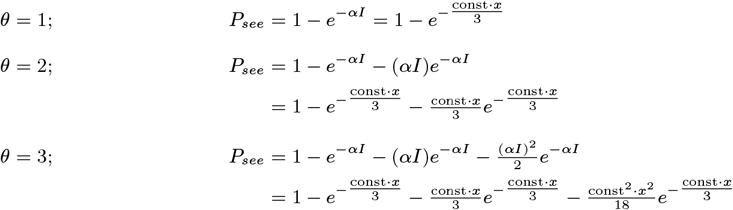

The fitted parameter in each case is *const*, a correction factor that accounts for photon dissipation and efficiency in signal transductions. For our experiments the expected value of *const* is close to 1 since photon loss at the the glass interfaces should be minimal, and we specifically target the end reaction of the signaling pathway.

#### Criteria for establishing independence of motor responses

We empirically determined the shortest interval between pulses for a specific blue light intensity (*I* = *I*_*min*_ · *x, x* = {1, 2, 4, 8, 16, 32}, *I*_*min*_ delivers ~ 7 · 10^12^ blue photons/cm^2^) that satisfied our criteria for non-interference (outlined below). For Opto-CheY_N449S_ (fast variant, ‘lit’ state half-life ≤ 1 sec) pulses are delivered every 2 seconds for *x* = 1, 2, every 3 seconds for *x* = 4, every 4 seconds for *x* = 8, every 5 seconds for *x* = 16, 32. For Opto-CheY_V416T_ (slow variant, ‘lit’ state half-life ≃ 2sec) pulses are delivered every 4 seconds for *x* = 1, every 6 seconds for *x* = 2, every 8 seconds for *x* = 4, every 10 seconds for *x* = 8, every 12 seconds for *x* = 16, every 15 seconds for *x* = 32. Below we outline how we selected these intervals:

- At low blue light intensities (low probability of motor response), we made an initial guess for pulse separation (3 seconds for the fast Opto-CheY), then reduced the separation to 2 seconds, and then to 1 second. The probability of motor response was similar for the 2- and 3-second intervals but increased significantly for the 1-second separation, suggesting that responses are not independent when pulses are applied every 1 second even at the lowest blue light intensity. To demonstrate this on the same cell in a least biased way, we performed the following experiment: low intensity pulses (equivalent to *x* = 4 intensity) are delivered in the following patterns:

1. pulse doublets (1 second apart) every 5 seconds.
2. pulse doublets (3 seconds apart) every 8 seconds.

Therefore, in the pulse sequence, the odd pulses are separated by 5 or 8 seconds from a previous pulse, while the even pulses are separated by 1 or 3 seconds from a previous pulse. If the response to the even pulse is conditioned by the odd pulse, the measured response probability for the even pulse should be markedly different from the response probability of the odd pulse (P_even_*>*P_odd_ under the assumption that the motor does not recover its baseline, and a higher baseline increases the response probability). Fig. S5 shows the CW bias of a motor that has undergone a pattern 1 followed by a pattern 2 stimulation (103 doublets of pattern 1, and 60 doublets of pattern 2). The probability of motor response was determined for the odd numbered pulses and, respectively, for the even numbered pulses. For pattern 1, P_odd_=0.087*±*0.028, while P_even_=0.165*±*0.037; for pattern 2, P_odd_=0.083*±*0.036 and P_even_=0.067*±*0.032. This experiment suggests that pulses separated by 3, 5 and 8 seconds generate independent motor responses, in contrast to pulses separated by 1 second.

- At higher blue light intensities the probability of motor response approaches saturation. We adjust the interval between pulses such that the motor bias returns to pre-stimulus values before the next stimulation. To demonstrate, we apply a pattern 1 stimulation (pulse doublets, 1 second apart, every 5 seconds) to individual cells/motors using higher intensity pulses (equivalent to the *x* = 16 pulses). Fig. S6 shows results from two different cells (a, b, c), and (a’, b’, c’). For both cells the ‘even’ stimulations happen before the motor has recovered its initial bias. For the first cell, differences in the response (first CW interval duration) are not evident at this sample size (58 doublet stimulations). The second cell, however, displays differences between the ‘odd’ and ‘even’ pulses both in the mean duration of the first CW interval as well as in its distribution. We conclude that return to initial bias in between stimulations is a stricter criterion.

### Supplementary Tables

**Table S1.**
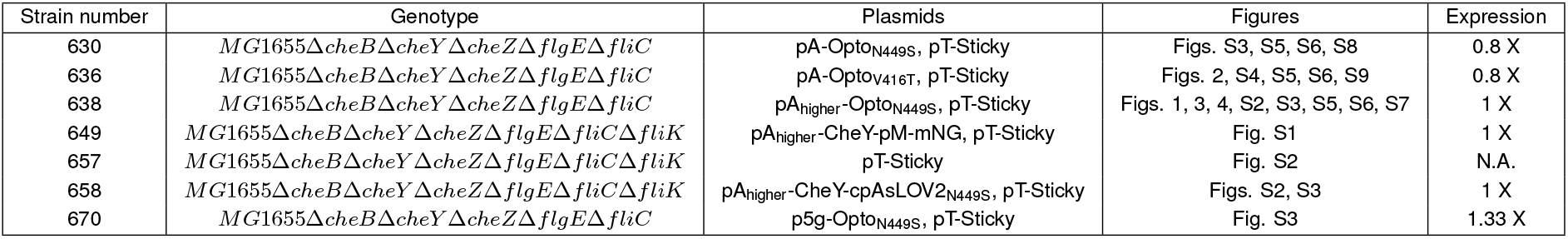
Experimental strains. The *E. coli* strains cited in the paper are described in terms of genotype and transformed plasmids. Column 4 shows the figures that show data from a particular strain. The last column displays the relative expression level of the Opto-CheY variant.

**Table S2.**
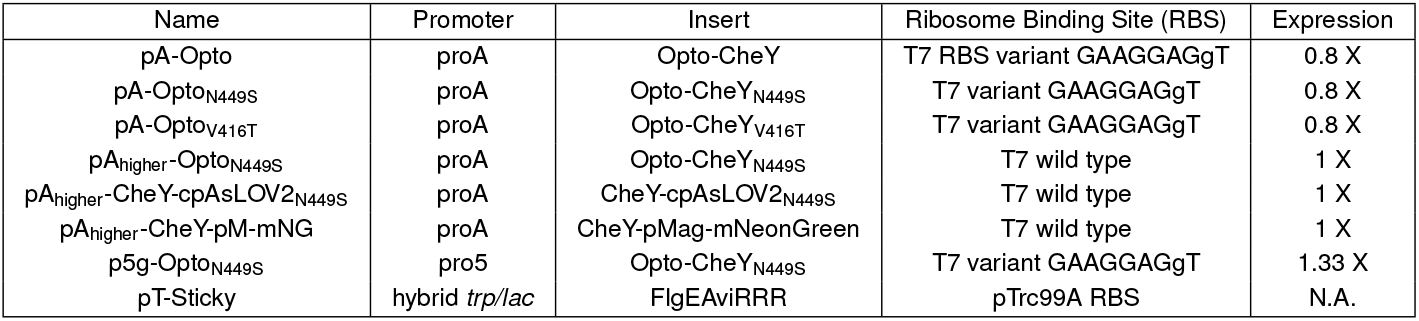
Plasmids. Transformed plasmids are described in terms of promoter variant, insert, and ribosome binding site (RBS). The combination of promoter strength and RBS variant gives the relative expression level for Opto-CheY.

**Table S3.**
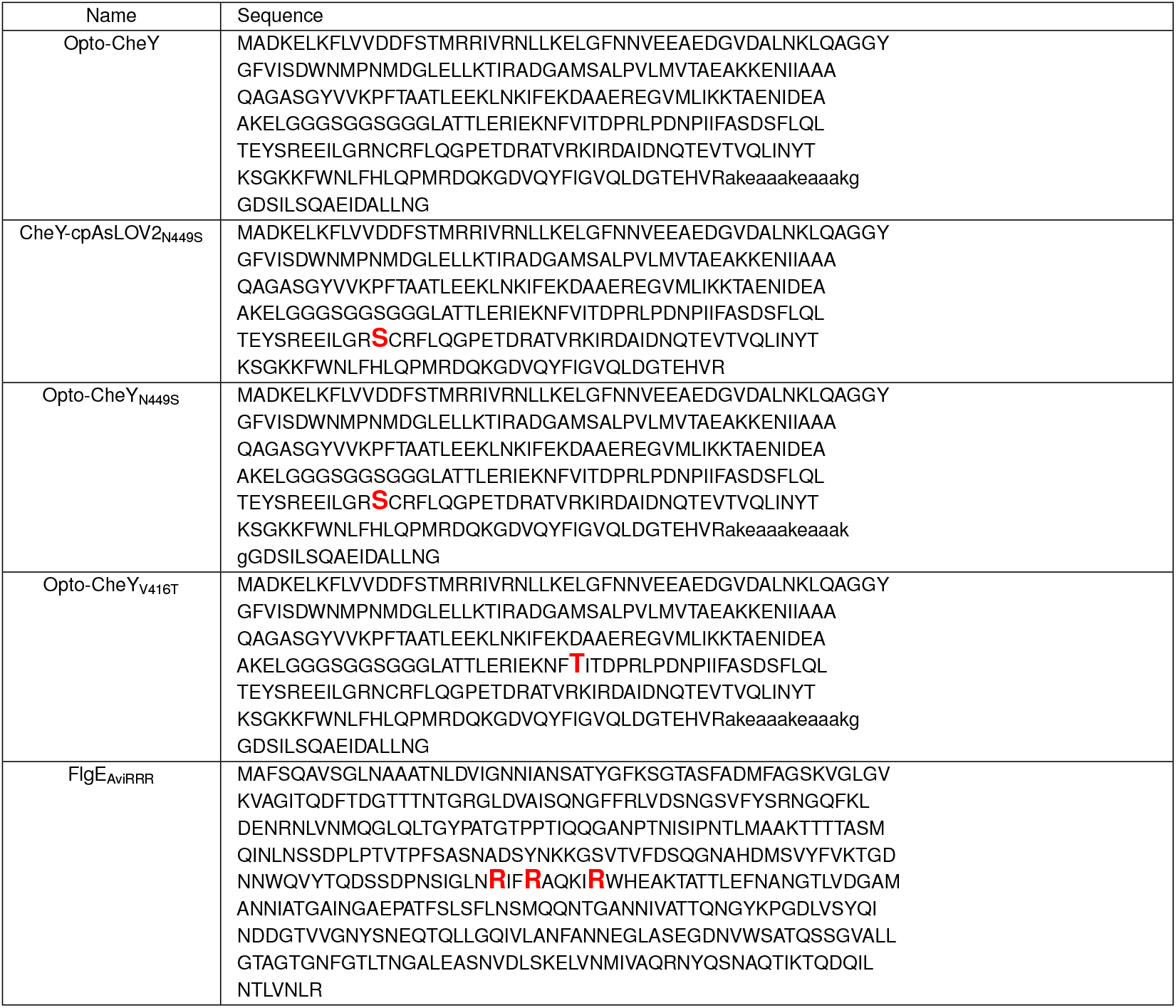
Sequences. The sequences of engineered proteins and variants are shown below. Red bold fonts highlight point mutations that were introduced in the wild type sequence.

**Table S4.**
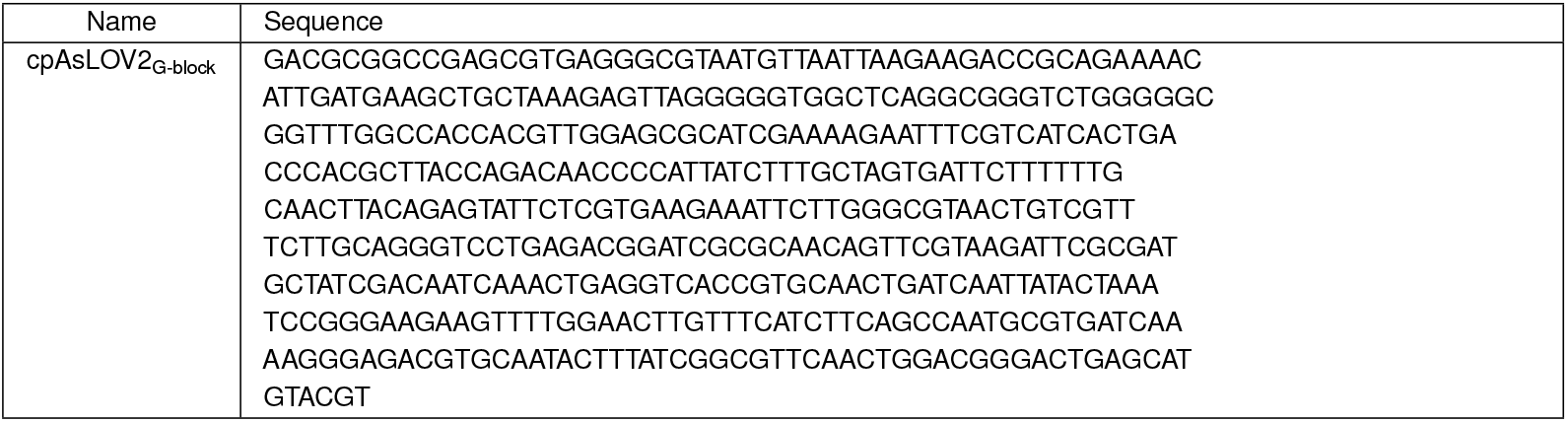
Nucleotide sequence of the cpAsLOV2 G-block synthesize by Invitrogen.

**Table S5.**
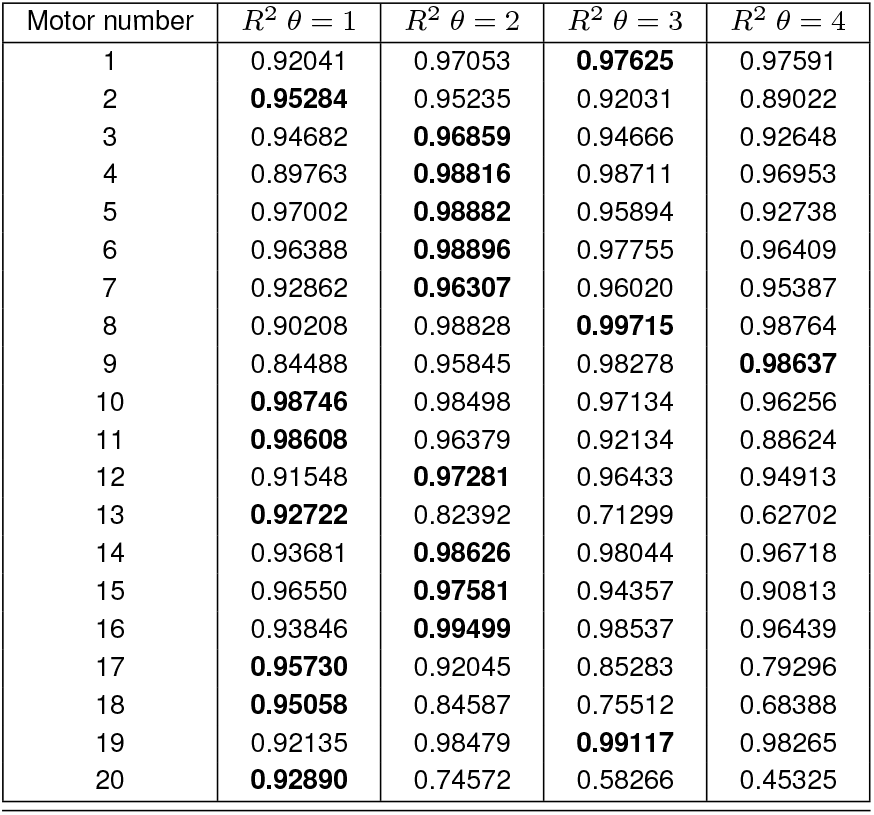
*R*^2^ values. Response probabilities for individual motors (1–20) of strain 636 were fitted to cumulative Poisson functions Psee with *θ* = 1 − 4. *R*^2^ values for each fit are shown with the highest value for a given motor in bold font. Note that these motors are also shown in Fig. 2 and Fig. S8 (V416T panel) and are numbered as an increasing function of response probability at the lowest blue stimulation.

### Supplemental Figures

**Fig. S1.**
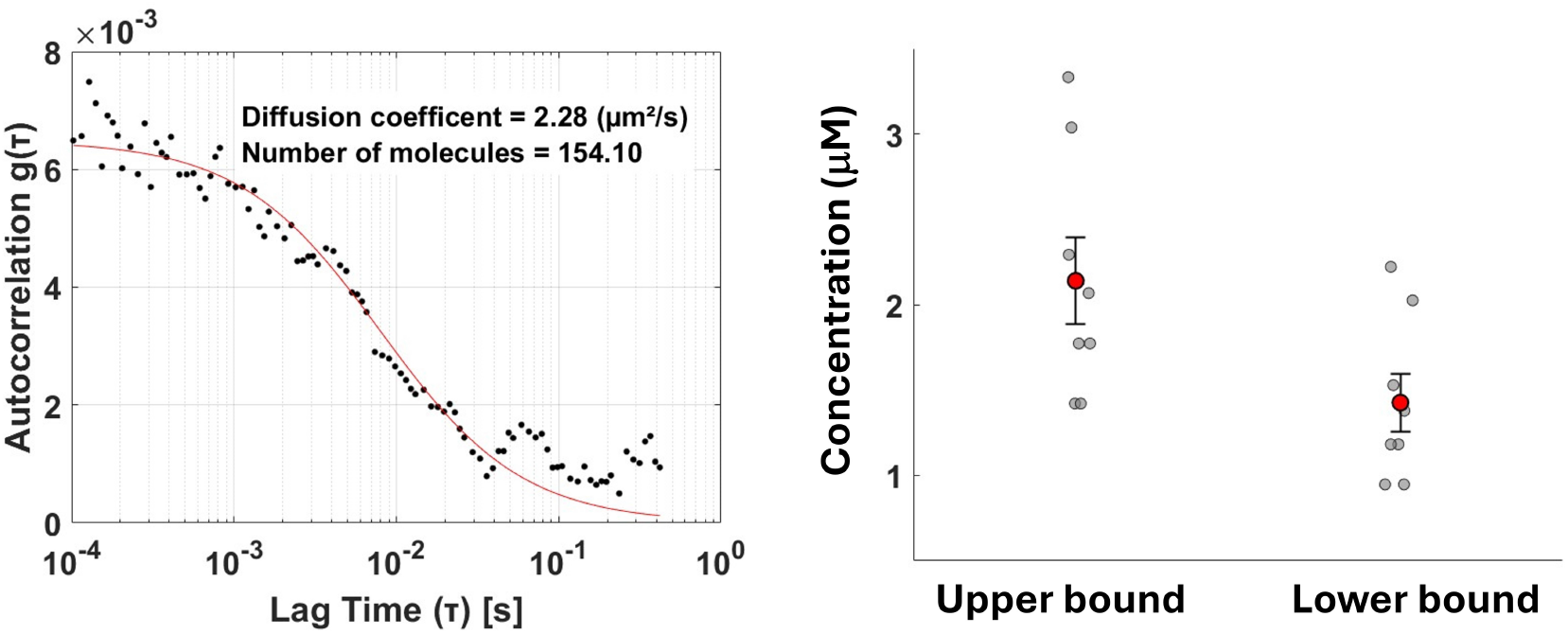
FCS for single bacteria. Left. Fluorescent signal autocorrelation (black dots) as a function of lag time for a bacterial cell. Data is fitted with a single component diffusion model (red line) with 2 parameters, diffusion time (used to compute the diffusion coefficient) and average number of molecules in the observed volume. **Right** Scatter plot for the computed upper bound and lower bound concentration for 8 cells (individual cell measurements are shown as gray circles). Mean upper bound is 2.14 *±* 0.25 µM and mean lower bound is 1.43 *±* 0.17 µM. Assuming the average volume of an *E. coli* bacterium to be 1 femto-L, each cell contains ~1000-2000 molecules of fluorescent protein.

**Fig. S2.**
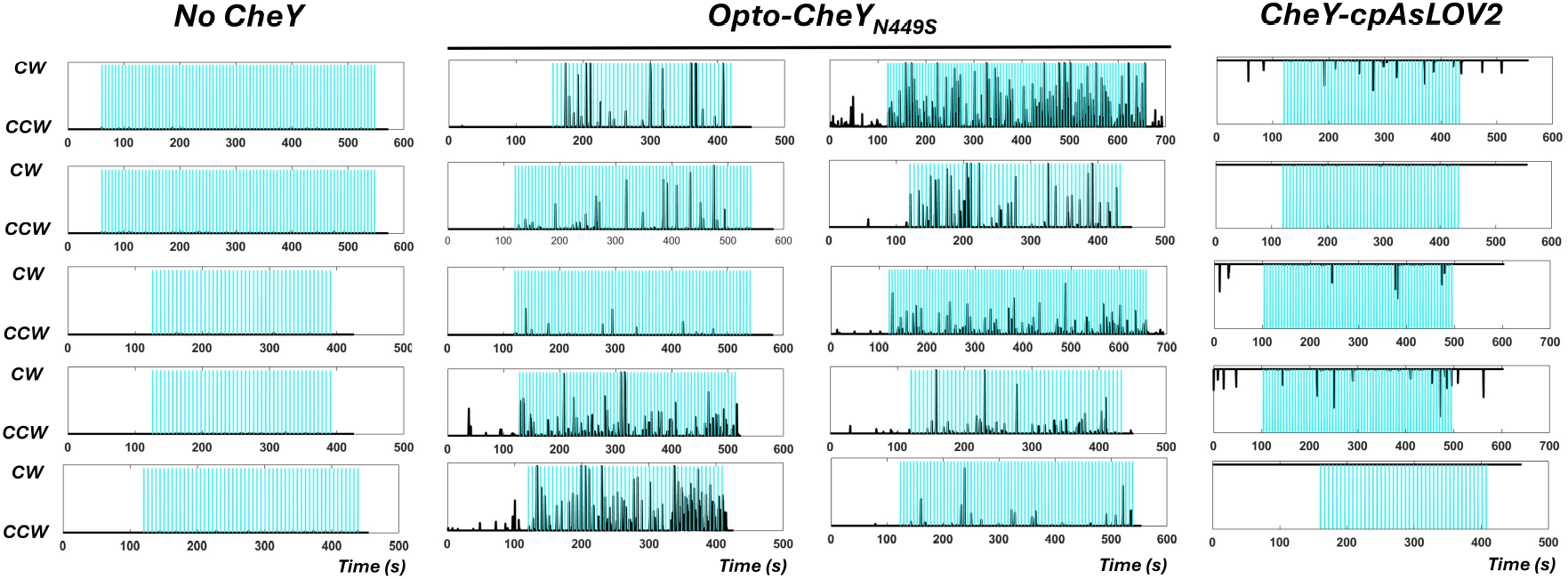
High intensity blue light pulses. Blue light pulses (2 msec, ~100 *µ*J/cm^2^) are applied every 6 seconds. Results are shown from 3 different strains: 657 (no plasmid), 638 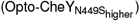, and 658 (CheY-cpAsLOV2_N449S_). Binary traces of cell body rotation (CCW=0, CW=1) are shown as a running average of 100 frames (500 msec). Blue light pulses are shown as vertical blue lines.

**Fig. S3.**
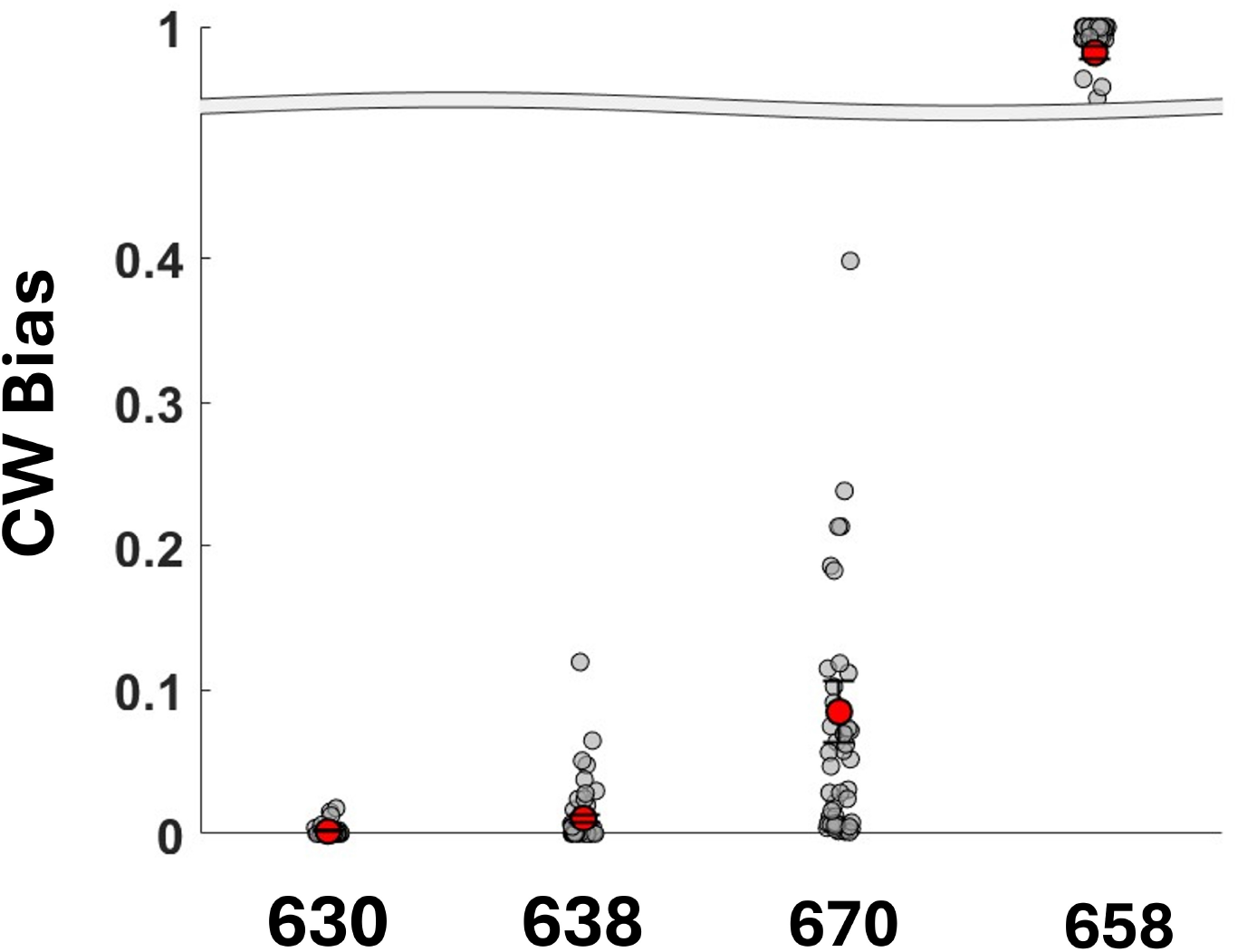
Initial CW bias. Scatter plot of initial CW biases for strains 630, 638, 670, and 658 (Table S1). Note the broken Y-axis. Individual measurements (gray), and means (red) with standard errors are shown. Mean CW biases are 0.00185 *±* 0.00064, 0.011 *±* 0.0032, 0.085 *±* 0.022 and 0.9821 *±* 0.0035, respectively.

**Fig. S4.**
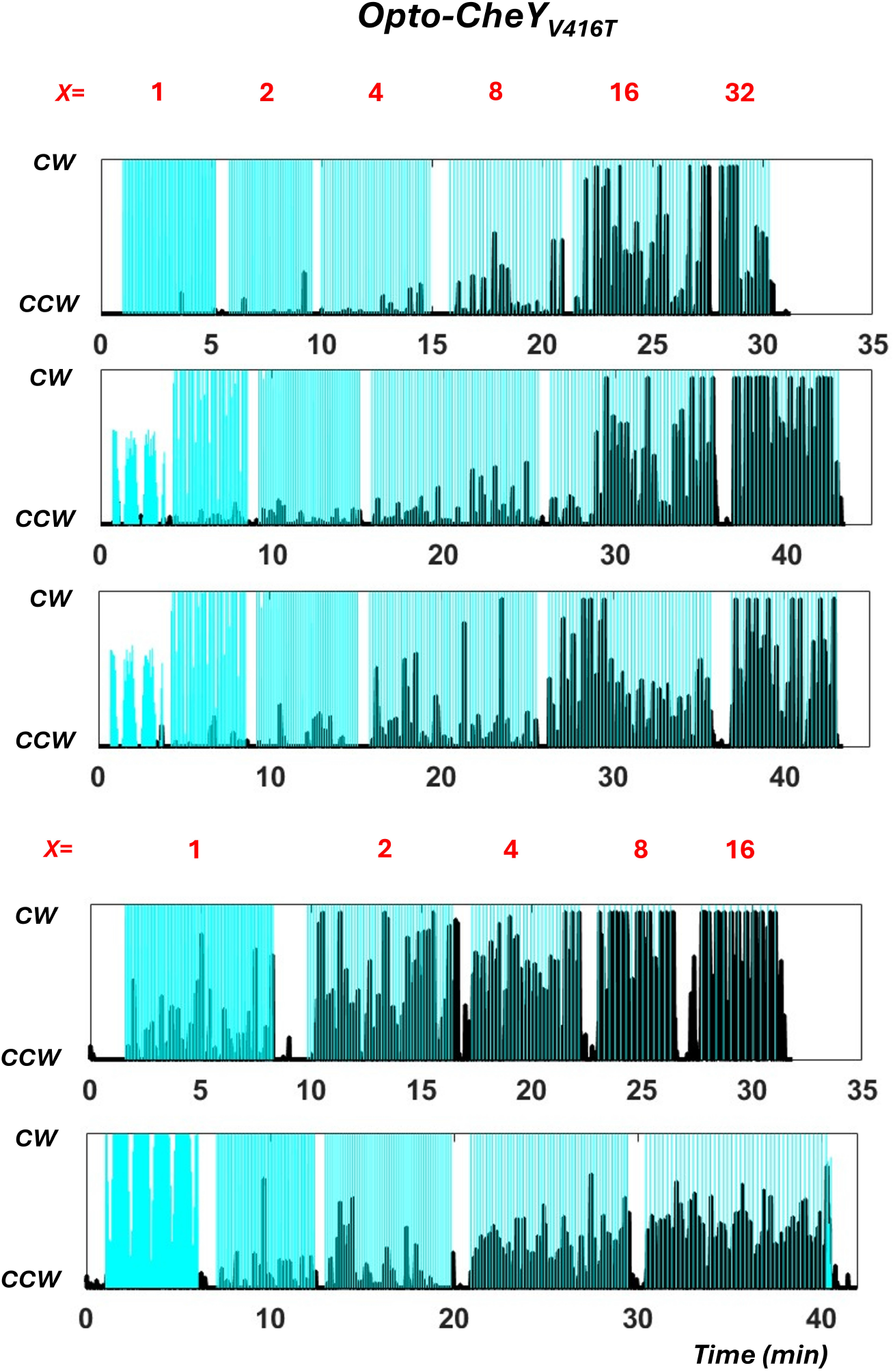
Motor responses to light pulses. Averaged binary traces (100 frames running average, 500 msec) for strain 636 (Opto-CheY_V416T_). Each plot is a single motor trace. Blue light pulses are shown as blue lines. 2 msec blue light pulses of a given blue light intensity (*I* = *I*_*min*_ · *x, x* = {1, 2, 4, 8, 16, 32}) are applied at constant frequency: every 4 sec for *x* = 1, every 6 sec for *x* = 2, every 8 sec for *x* = 4, every 10 sec for *x* = 8, every 12 sec for *x* = 16, and every 15 sec for *x* = 32.

**Fig. S5.**
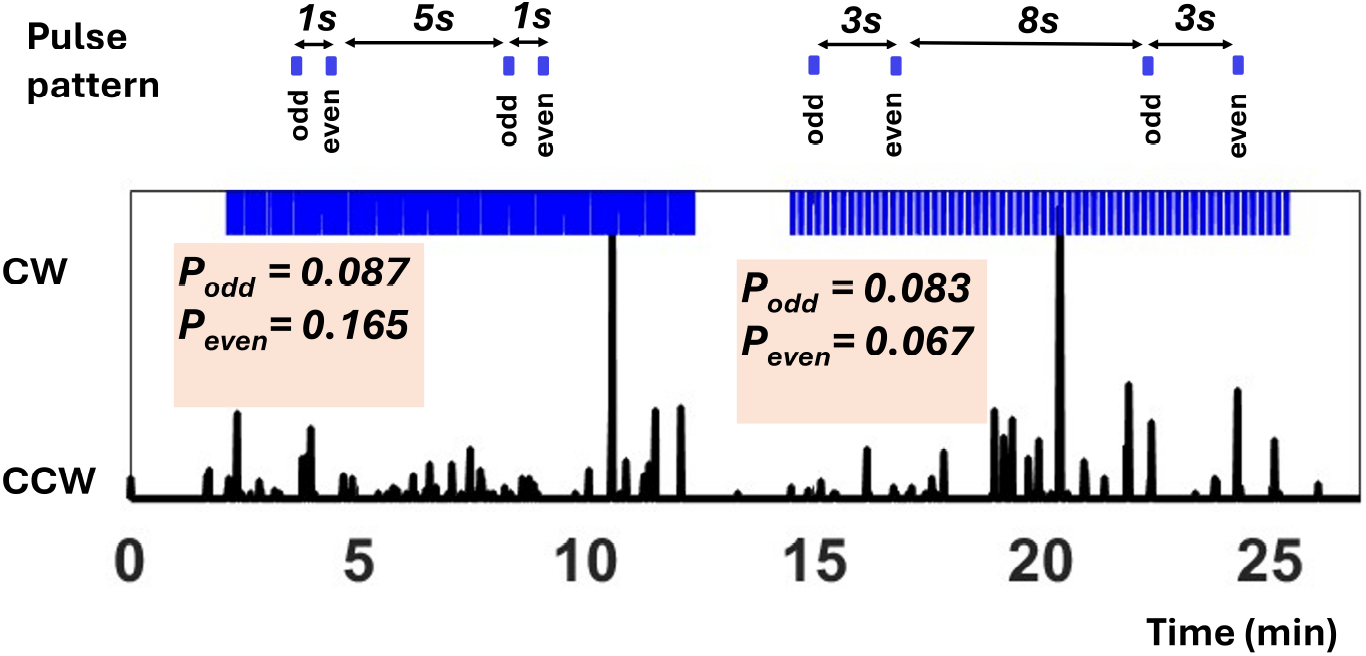
Response independence. Blue light pulses (*x* = 4) are delivered in patterns indicated above the individual motor trace (CW bias averaged with a 500 msec window). 103 doublets of pattern 1, and 60 doublets of pattern 2 were delivered. The probability of motor response was determined for the odd numbered pulses and, respectively, for the even numbered pulses. For pattern 1, P_odd_=0.087, while P_even_=0.165; for pattern 2, P_odd_=0.083 and P_even_=0.067.

**Fig. S6.**
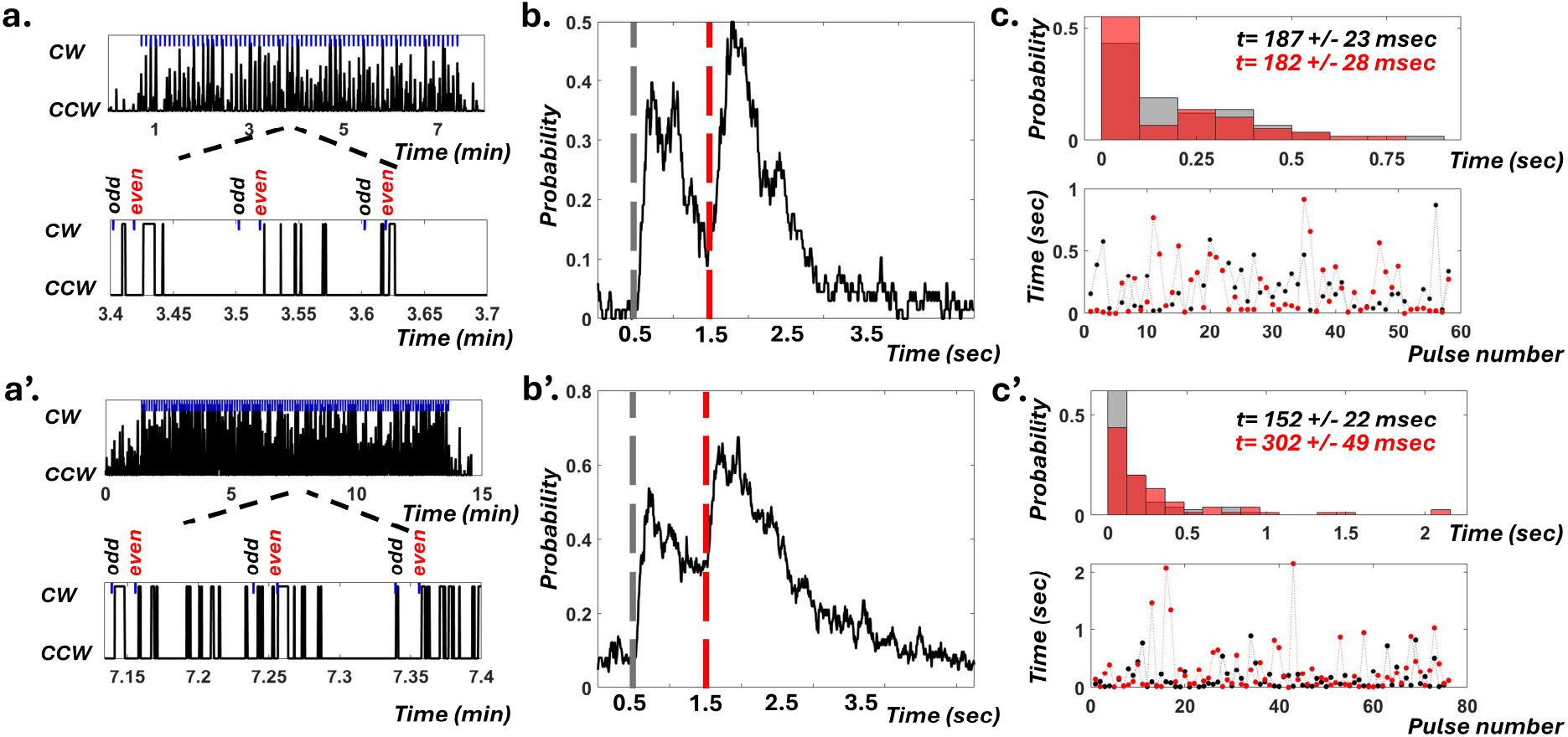
This figure shows data from two different cells (a, b, c), and (a’, b’, c’) with different initial CW average biases (0.03 and 0.06). Panels a/a’ show the cell’s CW bias as a function of time (100 frames (500 msec) running average for top panel and not averaged for the bottom zoom in). Blue light pulses are marked in blue. Panels b/b’ align and average all doublet stimulations giving the probability of motor response for the doublet stimulation as a function of time. The odd pulses are marked in gray, and the even pulses are marked in red. Note that odd pulses happen when the initial CW bias is recovered, while the even pulses happen when the motor’s bias is higher than the initial bias. Panels c/c’ show histograms for the duration of first CW interval following the odd pulses (gray), and even pulses (red). Average durations are also shown, as well as a plot of the first CW duration as a function of doublet pulse number (successive odd (gray) and even (red) pulses have the same doublet number).

**Fig. S7.**
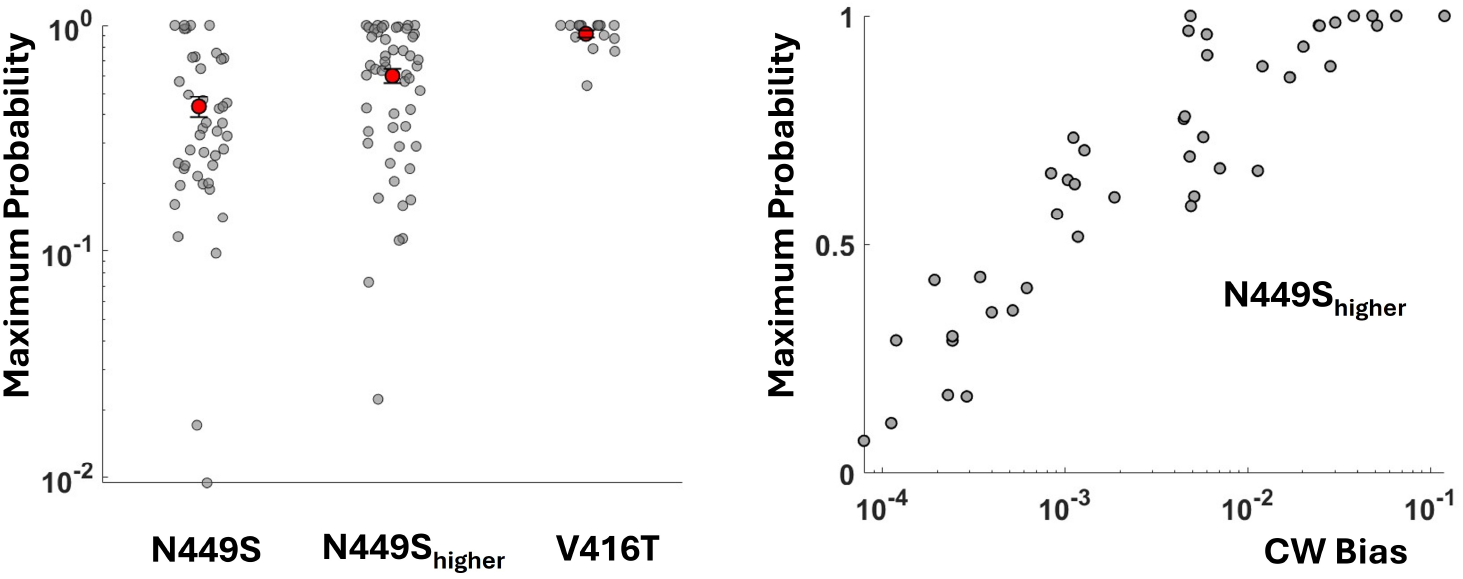
Saturating probability of the motor response. Left. Scatter plot of the maximal probability of motor response for Opto-Che 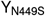, Opto-Che 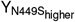, and Opto-Che 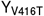motor (note logarithmic scale). Individual measurements (gray), and means (red) with standard errors are shown. Mean saturating probabilities are 0.435 *±* 0.0435, 0.6 *±* 0.0435 0.0437, and 0.92 *±* 0.034, respectively. **Right**. Maximum response probability as a function of initial CW bias for 51 Opto-Che 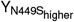motors.

**Fig. S8.**
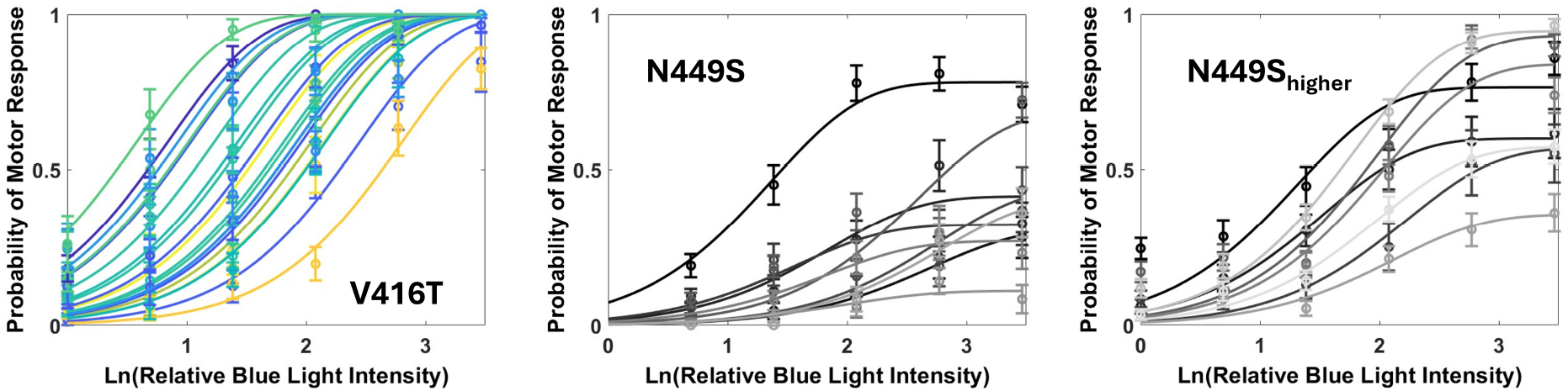
Blue light dose response curves for Opto-CheY variants. Opto-CheY_V416T_, Opto-CheY_N449S_, and Opto-Che 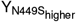 single motor dose response curves, as indicated. Each motor responds to light pulses of increasing relative intensity (*I* = *I*_*min*_ · *x, x* = {1, 2, 4, 8, 16, 32}) controlled using neutral density filters. Response probability for each BFM expressing an Opto-CheY variant at a given blue light intensity is calculated as the number of responses divided by the number of flashes. P_see_ with *θ* = 2 fits are shown for each of the 20 Opto-CheY_V416T_ dose response measurements in corresponding color. *R*^2^ values for the fits are given in Table S5. For Opto-CheY_N449S_ strains, data is fitted to *b×* P_see_, where *b* is between 0 and 1, a free parameter to allow response probability saturation at values less than 1.

**Fig. S9.**
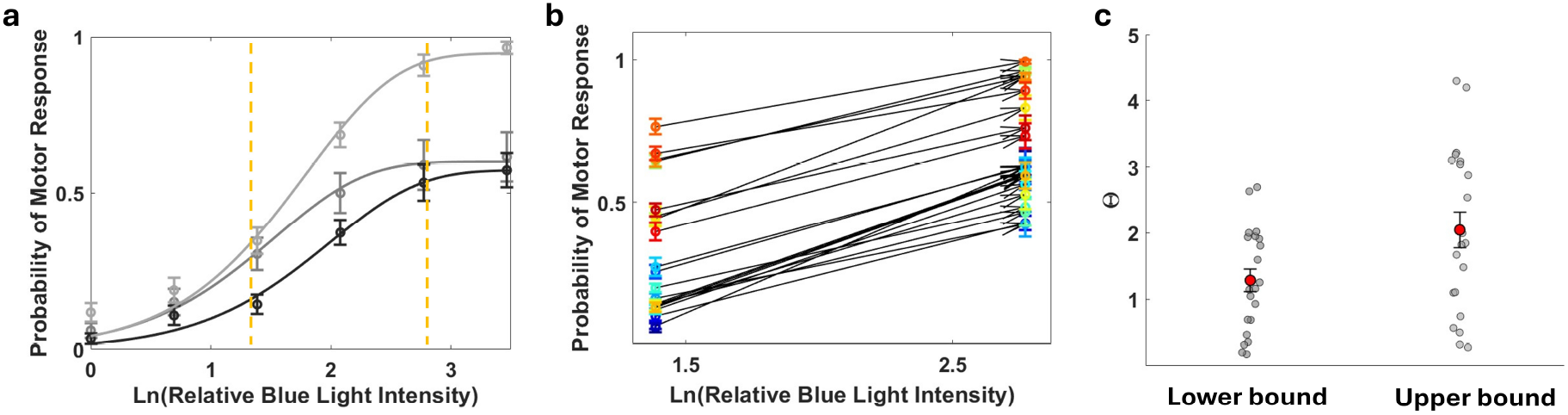
Response analysis for Opto-Che 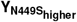. **a**. The dose response for three individual Opto-Che 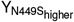motors is shown with corresponding *θ* = 2 fits. Vertical orange lines highlight the intensity interval with most rise in response probability (*x* = 4 − 16). **b**. 25 motors were stimulated at *x* = 4 (~ 300 flashes) and *x* = 16 (~ 100 flashes) intensities with enough trials to improve on response probability estimates. For each motor, the response probability estimates are connected by an arrow pointing towards the response probability for *x* = 16. **c**. *θ* can be obtained from the slope of the response as 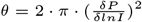, derivative evaluated at half maximal response. For each motor we can calculate the slope of the line connecting the *x* = 4 and *x* = 16 response probabilities. **c**. The lower bound *θ* estimate is made under the assumption that the response is approximately linear in the *x* = 4 − 16 interval, as it is the case for the black colored response in panel **a**. The upper bound estimate is for linear response in the *x* = 4 − 8 interval followed by saturation, as it is the case of the middle gray response in panel **a**. The upper bound estimate is for a linear response in the *x* = 4 − 8 interval followed by saturation, as it is the case of the middle gray response in panel **a**. *θ* estimates and corresponding initial CW biases of the motors are anti correlated, the higher the bias, the lower the *θ* estimate (Pearson correlation coefficient *r* = −0.7622, and *p* = 9.5038*e* − 06).

**Fig. S10.**
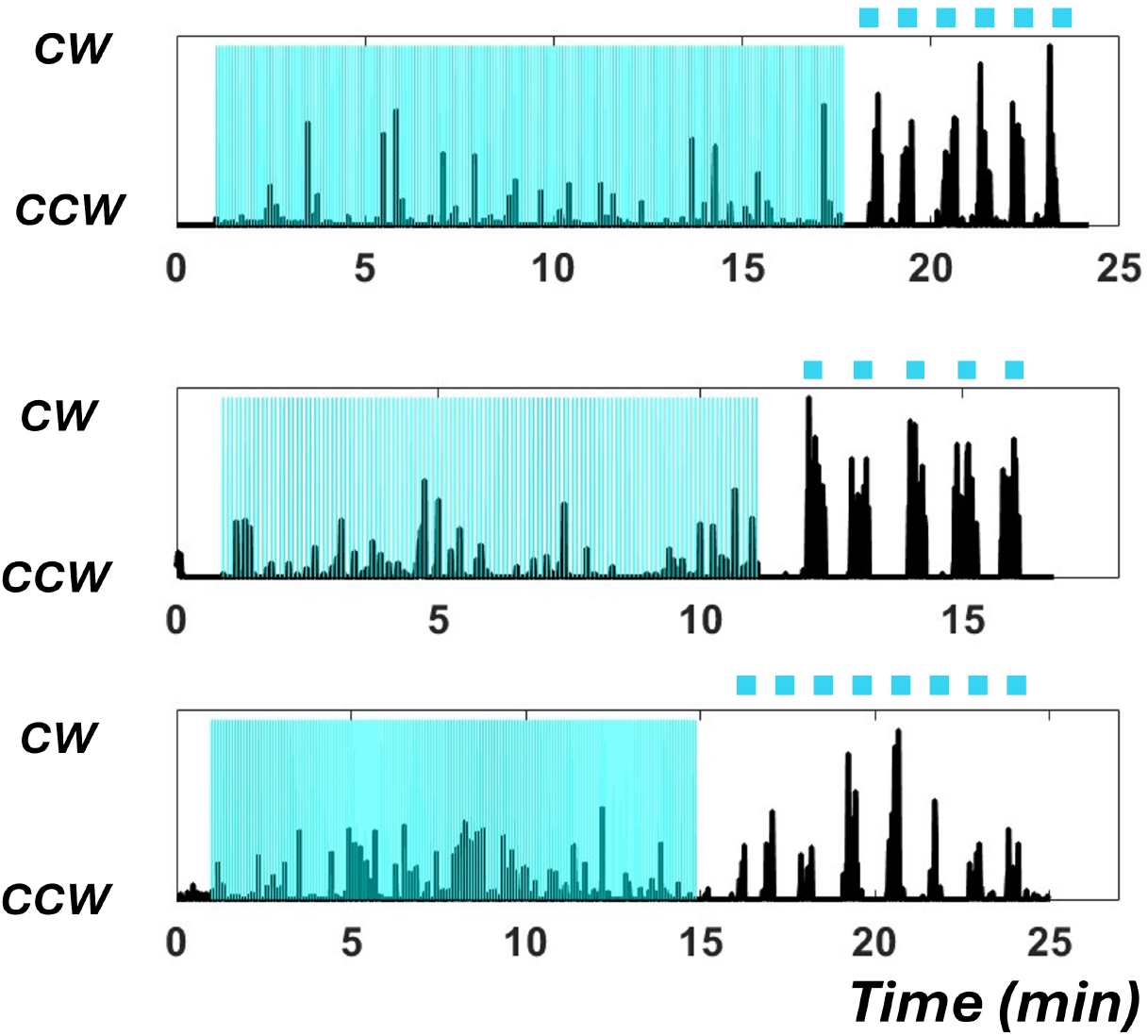
Longer pulses increase response probability. Averaged binary traces (100 frames running average, 500 msec) are shown for strain 630 (Table S1). Tethered cells (Opto-CheY_N449S_) are subjected to 2 msec pulses every 6 sec at the highest intensity (vertical blue lines). Following are 20 sec pulses (horizontal blue lines) at 1*/*32 of the maximum intensity.

**Fig. S11.**
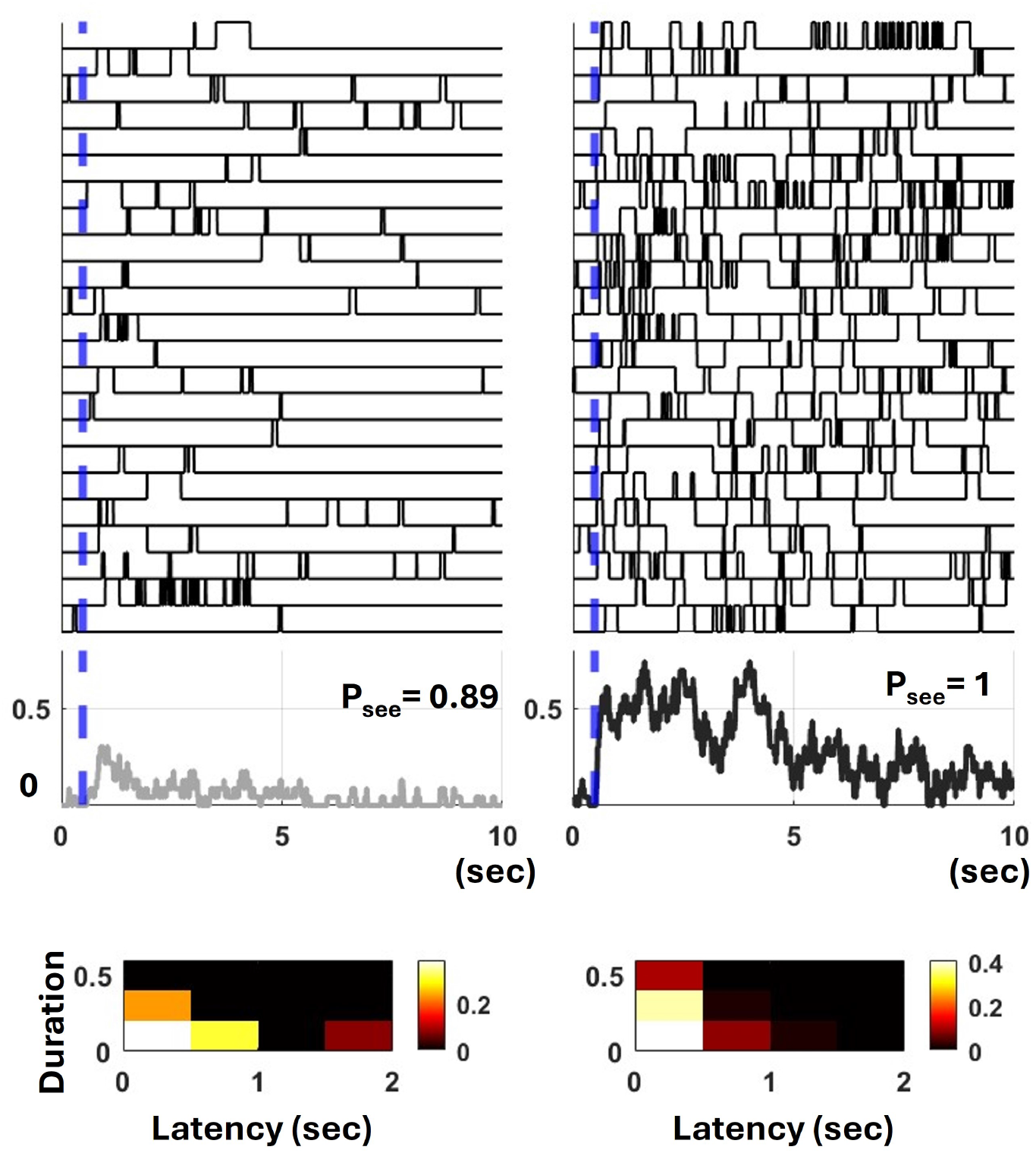
The CheY_V416T_ motor responses to different strength stimuli. A single motor is subjected to 28 light pulses (*I* = *I*_*min*_ · *x, x* = 4, weak stimulus), and 88 pulses (*x* = 16, strong stimulus) 23 responses are stacked and aligned relative to the light pulse (blue line). Average motor responses for each signal strength is shown (gray for the weak stimulus, and black for the strong stimulus). Below, the 23 latencies and durations are used to generate a joint probability distribution as a 2D-histogram, and color coded for probability as shown in the colorbar.

**Fig. S12.**
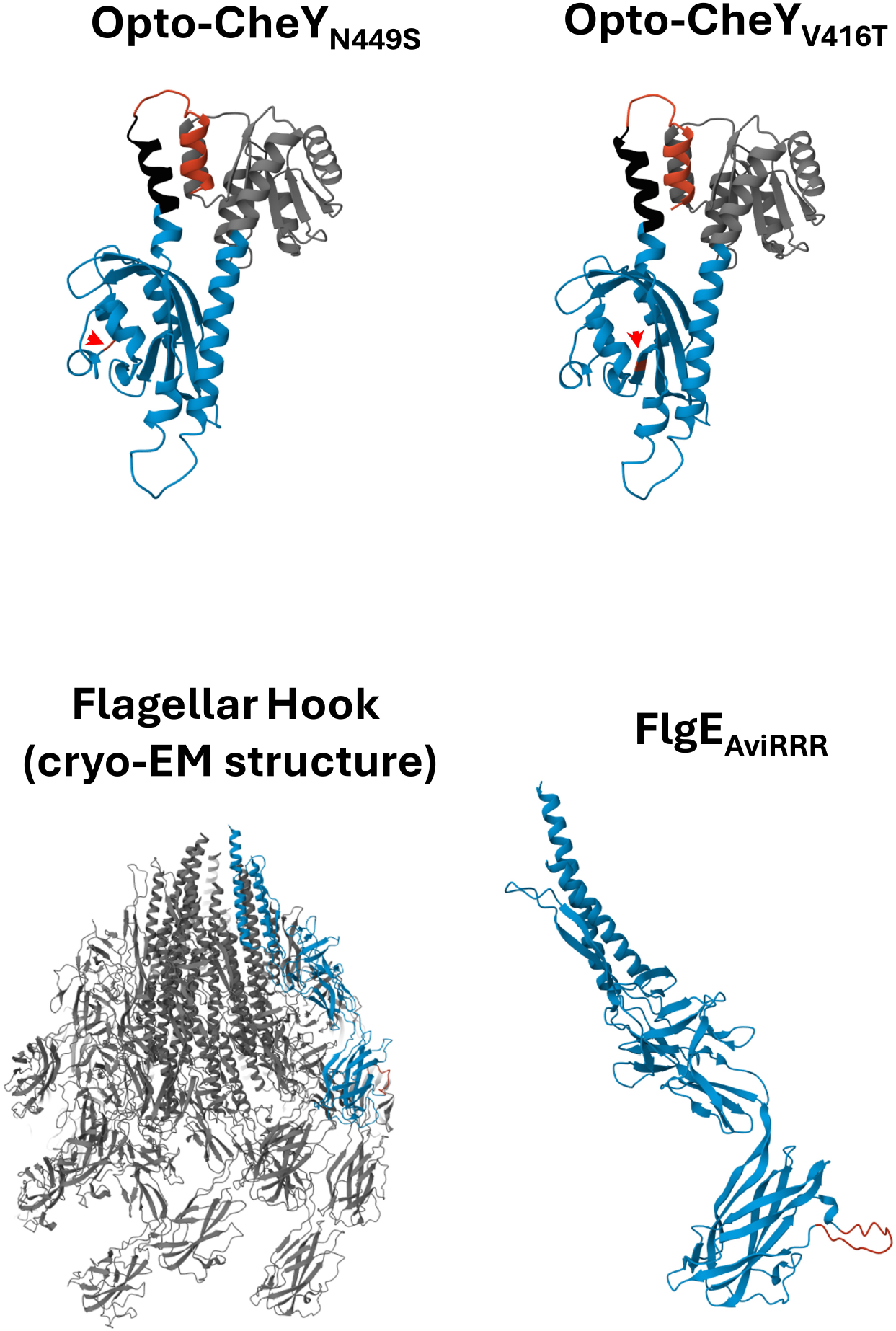
Structure predictions for engineered proteins. AlphaFold first ranked model predictions are shown for Opto-CheY_N449S_ and Opto-CheY_V416T_, color scheme as in Fig. 1b. Red arrow heads point to the location of the introduced mutations. Below, the cryo-EM structure of the flagellar hook from *Salmonella* is shown (pdb ID 7CGB).(4) One FlgE subunit of the hook is highlighted in blue. AlphaFold structure prediction of FlgE_AviRRR_ is shown to the right (engineered loop in red).

### Movies

Tethered cells movies are shown for strains 630, 636, 638, 657, 658 (Table S1). Each video shows 5 responses for the same cell, vertically stacked and aligned relative to the blue light pulse (2 msec, frame 101). For strain 636, blue light intensity is at 1*/*2 of the maximal intensity (~50 *µ*J/cm^2^, every 12 seconds (2400 frames)), while for the rest blue light intensity is maximal (~100 *µ*J/cm^2^, every 6 sec (1200 frames)). Videos are acquired at 200 fps, but shown here at 40 fps. A synchronized plot that tracks the rotational direction of the cell as a function of frame number is shown next to each video.

Movie 1. A 657 tethered cell.

Movie 2. A 638 tethered cell.

Movie 3. Another 638 tethered cell.

Movie 4. A 658 tethered cell.

Movie 5. A 630 tethered cell.

Movie 6. Another 630 tethered cell.

Movie 7. 636 tethered cell.

Movie 8. Another 636 tethered cell.

