## Supplementary Information Appendix for "The dynamic response of the bacterial flagellar motor to its direct intracellular input signal"

**Alina M. Vrabioiu, Basarab G. Hosu and Aravinthan D. T. Samuel (complete author list)**

**Corresponding Author name.**
****

**This PDF file includes:**

Supporting text
Figs. S1 to S12
Tables S1 to S5
SI References

### Supporting Information Text

### Supplementary methods

**Fluorescence correlation spectroscopy (FCS).** MG1655 $\Delta cheB \Delta cheY \Delta cheZ \Delta flgE \Delta fliC \Delta fliK$ cells express a CheY-pMag-mNeonGreen fusion using the proA promoter (strain number 649, Table S1). The plasmid backbone has kanamycin resistance and the low-copy origin SC101. (1) pMag is a variant of the fungal VVD photoresceptor of similar size and fold to AsLOV2 (2), and mNeonGreen was chosen for its increased photo-stability and fast maturation time.(1, 3) Cells were grown at 27$^o$ C till $OD_{600}$=0.6, and incubated at 10$^o$ C to ensure full maturation of the reporting fluorescent protein. Cells were then washed in motility buffer and attached to the poly lysine treated coverslip of a 35 mm MatTek dish. Zeiss LSM 880 confocal (40X water objective, $NA = 1.1$) and the Zeiss FCS software were used to acquire data at the Harvard Center for Biological Imaging. Alexa488 at a concentration of 30 nM was used to calibrate the system. We analyzed only the cells with $\leq 15\%$ fluorescent intensity decrease during the 2 sec acquisition time. We determined the average number of fluorescent molecules in the measured *E. coli* volume (Fig. S1). The width of *E. coli* is smaller than the axial dimension of the confocal volume ($\sim 844$ nm with confocal pinhole at 0.66 airy units). To convert to concentration, we considered a lower (500 nm), and upper (750 nm) bound for the width of *E. coli* (Fig. S1). Bleaching of mNeonGreen results in a slight underestimate of the number of molecules in the measured volume and of the CheY concentration.

**Blue light illumination.** The blue light LED was mounted on a side port on the Nikon Ti-U microscope, with neutral density filters that could be slided in and out of the light path (NDx, reduced light intensity by $1/x$). We measured the maximal LED power output used in our experiments to be $\sim 45\mu$W. For a 2 msec blue light pulse, that comes to $\sim 100nJ$ which corresponds to $\sim 2.2 \cdot 10^{11}$ blue light photons. The field of view (FOV) for our objective has a diameter of $\sim 400\mu$m and an area of $\sim 10^{-3} cm^2$. This gives a maximal energy output of $\sim 2.2 \cdot 10^{14}$ photons/$cm^2$; the minimal output is 1/32 of that which comes to $7 \cdot 10^{12}$ photons/$cm^2$. The probability of CheY activation at maximal energy is $p = \text{const} \cdot \sigma \cdot \Theta \cdot F_{max}$. For $\Theta \sim 40\%$, $\sigma \simeq 5.4 \cdot 10^{-17}$ cm$^2$, and $const = 1$, $p_{max} \simeq 0.005 \ll 1$.

**Data Fitting.** Individual motor dose-response data was fitted to the cumulative Poisson probability functions ($P_{see}$), plotted below for $\theta = 1 - 3$, $a$ is the average number of activated Opto-CheYs.

$$\theta = 1; \qquad P_{see} = 1 - e^{-a}$$

$$\theta = 2; \qquad P_{see} = 1 - e^{-a} - ae^{-a}$$

$$\theta = 3; \qquad P_{see} = 1 - e^{-a} - ae^{-a} - \frac{a^2}{2}e^{-a}$$

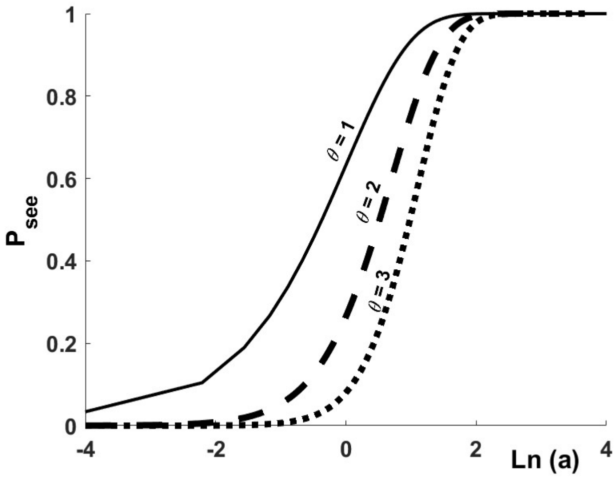


For our experiments $a = \alpha I = \text{const} \cdot N \cdot \sigma \cdot \Theta \cdot F_{min} \cdot x$, $1 \leq x \leq 32$. Since $F_{min} \sim 7 \cdot 10^{12}$ blue photons/cm$^2$, $N \sim 2000$ Opto-CheY molecules, $\Theta \sim 40\%$, and $\sigma \simeq 5.4 \cdot 10^{-17}$ cm$^2$, $\alpha I \simeq \frac{\text{const} \cdot x}{3}$.

$$\theta = 1; \qquad P_{see} = 1 - e^{-\alpha I} = 1 - e^{-\frac{\text{const}\cdot x}{3}}$$

$$\begin{aligned}\theta = 2; \qquad P_{see} &= 1 - e^{-\alpha I} - (\alpha I)e^{-\alpha I} \\ &= 1 - e^{-\frac{\text{const}\cdot x}{3}} - \frac{\text{const}\cdot x}{3} e^{-\frac{\text{const}\cdot x}{3}}\end{aligned}$$

$$\begin{aligned}\theta = 3; \qquad P_{see} &= 1 - e^{-\alpha I} - (\alpha I)e^{-\alpha I} - \frac{(\alpha I)^2}{2} e^{-\alpha I} \\ &= 1 - e^{-\frac{\text{const}\cdot x}{3}} - \frac{\text{const}\cdot x}{3} e^{-\frac{\text{const}\cdot x}{3}} - \frac{\text{const}^2\cdot x^2}{18} e^{-\frac{\text{const}\cdot x}{3}}\end{aligned}$$

The fitted parameter in each case is *const*, a correction factor that accounts for photon dissipation and efficiency in signal transductions. For our experiments the expected value of *const* is close to 1 since photon loss at the the glass interfaces should be minimal, and we specifically target the end reaction of the signaling pathway.

**Criteria for establishing independence of motor responses.** We empirically determined the shortest interval between pulses for a specific blue light intensity ($I = I_{min} \cdot x$, $x = \{1, 2, 4, 8, 16, 32\}$, $I_{min}$ delivers $\sim 7 \cdot 10^{12}$ blue photons/cm$^2$) that satisfied our criteria for non-interference (outlined below). For Opto-CheY$_{\text{N449S}}$ (fast variant, 'lit' state half-life $\leq$ 1 sec) pulses are delivered every 2 seconds for $x = 1, 2$, every 3 seconds for $x = 4$, every 4 seconds for $x = 8$, every 5 seconds for $x = 16, 32$. For Opto-CheY$_{\text{V416T}}$ (slow variant, 'lit' state half-life $\simeq$ 2sec) pulses are delivered every 4 seconds for $x = 1$, every 6 seconds for $x = 2$, every 8 seconds for $x = 4$, every 10 seconds for $x = 8$, every 12 seconds for $x = 16$, every 15 seconds for $x = 32$. Below we outline how we selected these intervals:

### Supplementary Tables

**Table S1. Experimental strains. The *E. coli* strains cited in the paper are described in terms of genotype and transformed plasmids. Column 4 shows the figures that show data from a particular strain. The last column displays the relative expression level of the Opto-CheY variant.**

| Strain number | Genotype | Plasmids | Figures | Expression |
|---|---|---|---|---|
| 630 | $MG1655\Delta cheB\Delta cheY\Delta cheZ\Delta flgE\Delta fliC$ | pA-Opto$_{N449S}$, pT-Sticky | Figs. S3, S5, S6, S8 | 0.8 X |
| 636 | $MG1655\Delta cheB\Delta cheY\Delta cheZ\Delta flgE\Delta fliC$ | pA-Opto$_{V416T}$, pT-Sticky | Figs. 2, S4, S5, S6, S9 | 0.8 X |
| 638 | $MG1655\Delta cheB\Delta cheY\Delta cheZ\Delta flgE\Delta fliC$ | pA$_{higher}$-Opto$_{N449S}$, pT-Sticky | Figs. 1, 3, 4, S2, S3, S5, S6, S7 | 1 X |
| 649 | $MG1655\Delta cheB\Delta cheY\Delta cheZ\Delta flgE\Delta fliC\Delta fliK$ | pA$_{higher}$-CheY-pM-mNG, pT-Sticky | Fig. S1 | 1 X |
| 657 | $MG1655\Delta cheB\Delta cheY\Delta cheZ\Delta flgE\Delta fliC\Delta fliK$ | pT-Sticky | Fig. S2 | N.A. |
| 658 | $MG1655\Delta cheB\Delta cheY\Delta cheZ\Delta flgE\Delta fliC\Delta fliK$ | pA$_{higher}$-CheY-cpAsLOV2$_{N449S}$, pT-Sticky | Figs. S2, S3 | 1 X |
| 670 | $MG1655\Delta cheB\Delta cheY\Delta cheZ\Delta flgE\Delta fliC$ | p5g-Opto$_{N449S}$, pT-Sticky | Fig. S3 | 1.33 X |

**Table S2. Plasmids. Transformed plasmids are described in terms of promoter variant, insert, and ribosome binding site (RBS). The combination of promoter strength and RBS variant gives the relative expression level for Opto-CheY.**

| Name | Promoter | Insert | Ribosome Binding Site (RBS) | Expression |
|---|---|---|---|---|
| pA-Opto | proA | Opto-CheY | T7 RBS variant GAAGGAGgT | 0.8 X |
| pA-Opto$_{N449S}$ | proA | Opto-CheY$_{N449S}$ | T7 variant GAAGGAGgT | 0.8 X |
| pA-Opto$_{V416T}$ | proA | Opto-CheY$_{V416T}$ | T7 variant GAAGGAGgT | 0.8 X |
| pA$_{higher}$-Opto$_{N449S}$ | proA | Opto-CheY$_{N449S}$ | T7 wild type | 1 X |
| pA$_{higher}$-CheY-cpAsLOV2$_{N449S}$ | proA | CheY-cpAsLOV2$_{N449S}$ | T7 wild type | 1 X |
| pA$_{higher}$-CheY-pM-mNG | proA | CheY-pMag-mNeonGreen | T7 wild type | 1 X |
| p5g-Opto$_{N449S}$ | pro5 | Opto-CheY$_{N449S}$ | T7 variant GAAGGAGgT | 1.33 X |
| pT-Sticky | hybrid *trp/lac* | FlgEAviRRR | pTrc99A RBS | N.A. |

**Table S3. Sequences. The sequences of engineered proteins and variants are shown below. Red bold fonts highlight point mutations that were introduced in the wild type sequence.**

| Name | Sequence |
|---|---|
| Opto-CheY | MADKELKFLVVDDFSTMRRIVRNLLKELGFNNVEEAEDGVDALNKLQAGGY<br>GFVISDWNMPNMDGLELLKTIRADGAMSALPVLMVTAEAKKENIIAAA<br>QAGASGYVVKPFTAATLEEKLNKIFEKDAAEREGVMLIKKTAENIDEA<br>AKELGGGSGGSGGGLATTLERIEKNFVITDPRLPDNPIIFASDSFLQL<br>TEYSREEILGRNCRFLQGPETDRATVRKIRDAIDNQTEVTVQLINYT<br>KSGKKFWNLFHLQPMRDQKGDVQYFIGVQLDGTEHVRakeaaakeaaakg<br>GDSILSQAEIDALLNG |
| CheY-cpAsLOV2$_{N449S}$ | MADKELKFLVVDDFSTMRRIVRNLLKELGFNNVEEAEDGVDALNKLQAGGY<br>GFVISDWNMPNMDGLELLKTIRADGAMSALPVLMVTAEAKKENIIAAA<br>QAGASGYVVKPFTAATLEEKLNKIFEKDAAEREGVMLIKKTAENIDEA<br>AKELGGGSGGSGGGLATTLERIEKNFVITDPRLPDNPIIFASDSFLQL<br>TEYSREEILGR**S**CRFLQGPETDRATVRKIRDAIDNQTEVTVQLINYT<br>KSGKKFWNLFHLQPMRDQKGDVQYFIGVQLDGTEHVR |
| Opto-CheY$_{N449S}$ | MADKELKFLVVDDFSTMRRIVRNLLKELGFNNVEEAEDGVDALNKLQAGGY<br>GFVISDWNMPNMDGLELLKTIRADGAMSALPVLMVTAEAKKENIIAAA<br>QAGASGYVVKPFTAATLEEKLNKIFEKDAAEREGVMLIKKTAENIDEA<br>AKELGGGSGGSGGGLATTLERIEKNFVITDPRLPDNPIIFASDSFLQL<br>TEYSREEILGR**S**CRFLQGPETDRATVRKIRDAIDNQTEVTVQLINYT<br>KSGKKFWNLFHLQPMRDQKGDVQYFIGVQLDGTEHVRakeaaakeaaak<br>gGDSILSQAEIDALLNG |
| Opto-CheY$_{V416T}$ | MADKELKFLVVDDFSTMRRIVRNLLKELGFNNVEEAEDGVDALNKLQAGGY<br>GFVISDWNMPNMDGLELLKTIRADGAMSALPVLMVTAEAKKENIIAAA<br>QAGASGYVVKPFTAATLEEKLNKIFEKDAAEREGVMLIKKTAENIDEA<br>AKELGGGSGGSGGGLATTLERIEKNF**T**ITDPRLPDNPIIFASDSFLQL<br>TEYSREEILGRNCRFLQGPETDRATVRKIRDAIDNQTEVTVQLINYT<br>KSGKKFWNLFHLQPMRDQKGDVQYFIGVQLDGTEHVRakeaaakeaaakg<br>GDSILSQAEIDALLNG |
| FlgE$_{AviRRR}$ | MAFSQAVSGLNAAATNLDVIGNNIANSATYGFKSGTASFADMFAGSKVGLGV<br>KVAGITQDFTDGTTTNTGRGLDVAISQNGFFRLVDSNGSVFYSRNGQFKL<br>DENRNLVNMQGLQLTGYPATGTPPTIQQGANPTNISIPNTLMAAKTTTTASM<br>QINLNSSDPLPTVTPFSASNADSYNKKGSVTVFDSQGNAHDMSVYFVKTGD<br>NNWQVYTQDSSDPNSIGLN**R**IF**R**AQKI**R**WHEAKTATTLEFNANGTLVDGAM<br>ANNIATGAINGAEPATFSLSFLNSMQQNTGANNIVATTQNGYKPGDLVSYQI<br>NDDGTVVGNYSNEQTQLLGQIVLANFANNEGLASEGDNVWSATQSSGVALL<br>GTAGTGNFGTLTNGALEASNVDLSKELVNMIVAQRNYQSNAQTIKTQDQIL<br>NTLVNLR |

**Table S4. Nucleotide sequence of the cpAsLOV2 G-block synthesize by Invitrogen.**

| Name | Sequence |
|---|---|
| cpAsLOV2$_{\text{G-block}}$ | GACGCGGCCGAGCGTGAGGGCGTAATGTTAATTAAGAAGACCGCAGAAAAC<br>ATTGATGAAGCTGCTAAAGAGTTAGGGGGTGGCTCAGGCGGGTCTGGGGGC<br>GGTTTGGCCACCACGTTGGAGCGCATCGAAAAGAATTTCGTCATCACTGA<br>CCCACGCTTACCAGACAACCCCATTATCTTTGCTAGTGATTCTTTTTTG<br>CAACTTACAGAGTATTCTCGTGAAGAAATTCTTGGGCGTAACTGTCGTT<br>TCTTGCAGGGTCCTGAGACGGATCGCGCAACAGTTCGTAAGATTCGCGAT<br>GCTATCGACAATCAAACTGAGGTCACCGTGCAACTGATCAATTATACTAAA<br>TCCGGGAAGAAGTTTTGGAACTTGTTTCATCTTCAGCCAATGCGTGATCAA<br>AAGGGAGACGTGCAATACTTTATCGGCGTTCAACTGGACGGGACTGAGCAT<br>GTACGT |

**Table S5.** $R^2$ **values. Response probabilities for individual motors (1-20) of strain 636 were fitted to cumulative Poisson functions $P_{see}$ with $\theta = 1 - 4$. $R^2$ values for each fit are shown with the highest value for a given motor in bold font. Note that these motors are also shown in Fig. 2 and Fig. S8 (V416T panel) and are numbered as an increasing function of response probability at the lowest blue stimulation.**

| Motor number | $R^2$ $\theta = 1$ | $R^2$ $\theta = 2$ | $R^2$ $\theta = 3$ | $R^2$ $\theta = 4$ |
|---|---|---|---|---|
| 1 | 0.92041 | 0.97053 | **0.97625** | 0.97591 |
| 2 | **0.95284** | 0.95235 | 0.92031 | 0.89022 |
| 3 | 0.94682 | **0.96859** | 0.94666 | 0.92648 |
| 4 | 0.89763 | **0.98816** | 0.98711 | 0.96953 |
| 5 | 0.97002 | **0.98882** | 0.95894 | 0.92738 |
| 6 | 0.96388 | **0.98896** | 0.97755 | 0.96409 |
| 7 | 0.92862 | **0.96307** | 0.96020 | 0.95387 |
| 8 | 0.90208 | 0.98828 | **0.99715** | 0.98764 |
| 9 | 0.84488 | 0.95845 | 0.98278 | **0.98637** |
| 10 | **0.98746** | 0.98498 | 0.97134 | 0.96256 |
| 11 | **0.98608** | 0.96379 | 0.92134 | 0.88624 |
| 12 | 0.91548 | **0.97281** | 0.96433 | 0.94913 |
| 13 | **0.92722** | 0.82392 | 0.71299 | 0.62702 |
| 14 | 0.93681 | **0.98626** | 0.98044 | 0.96718 |
| 15 | 0.96550 | **0.97581** | 0.94357 | 0.90813 |
| 16 | 0.93846 | **0.99499** | 0.98537 | 0.96439 |
| 17 | **0.95730** | 0.92045 | 0.85283 | 0.79296 |
| 18 | **0.95058** | 0.84587 | 0.75512 | 0.68388 |
| 19 | 0.92135 | 0.98479 | **0.99117** | 0.98265 |
| 20 | **0.92890** | 0.74572 | 0.58266 | 0.45325 |

### Supplemental Figures

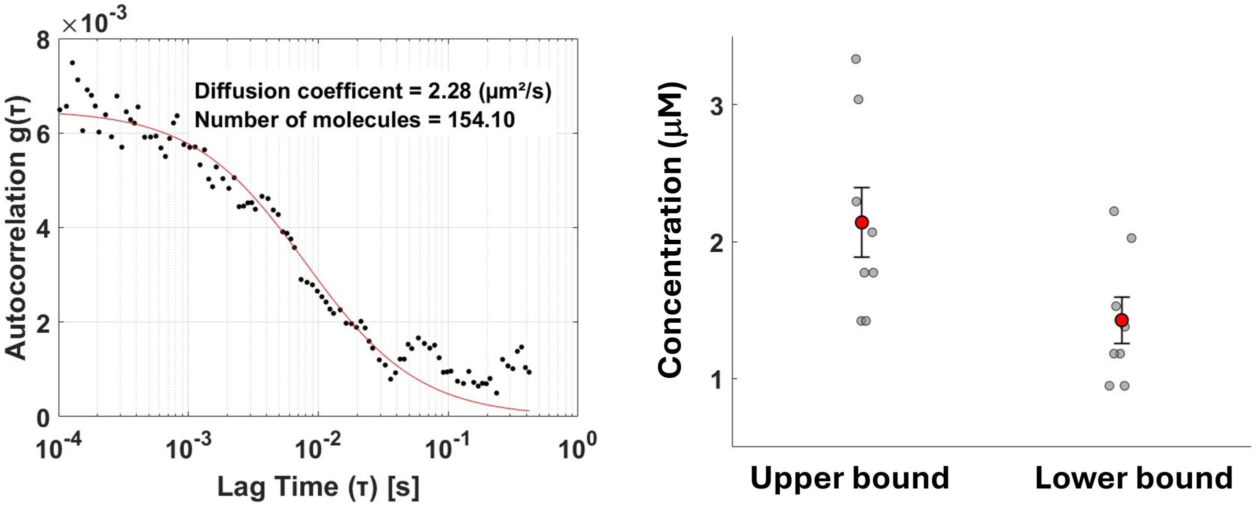


**Fig. S1. FCS for single bacteria. Left.** Fluorescent signal autocorrelation (black dots) as a function of lag time for a bacterial cell. Data is fitted with a single component diffusion model (red line) with 2 parameters, diffusion time (used to compute the diffusion coefficient) and average number of molecules in the observed volume. **Right** Scatter plot for the computed upper bound and lower bound concentration for 8 cells (individual cell measurements are shown as gray circles). Mean upper bound is $2.14 \pm 0.25$ µM and mean lower bound is $1.43 \pm 0.17$ µM. Assuming the average volume of an *E. coli* bacterium to be 1 femto-L, each cell contains ~1000-2000 molecules of fluorescent protein.

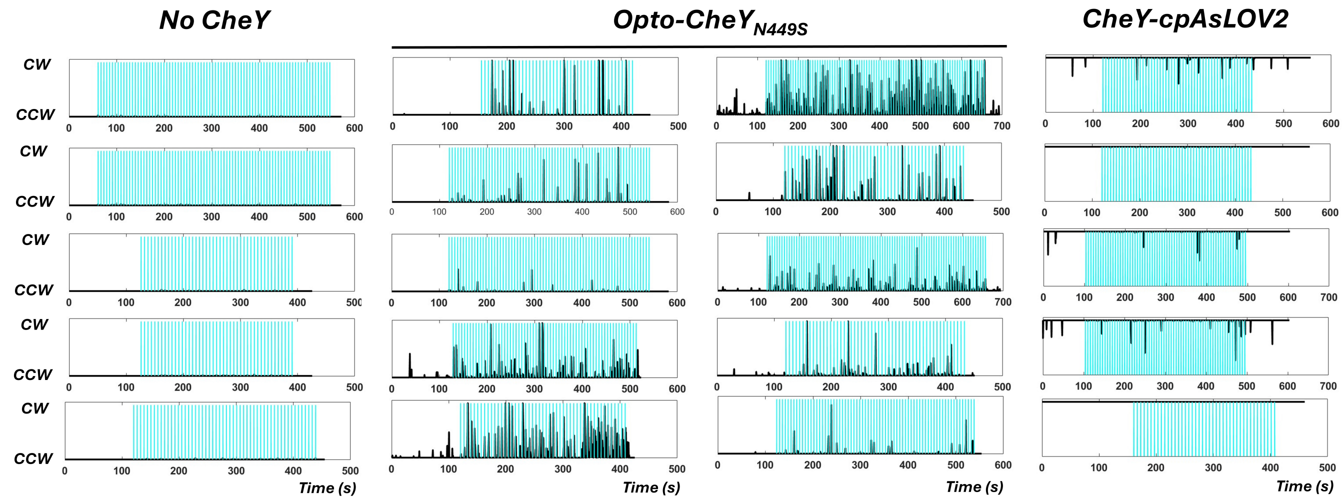


**Fig. S2. High intensity blue light pulses.** Blue light pulses (2 msec, ~100 $\mu$J/cm$^2$) are applied every 6 seconds. Results are shown from 3 different strains: 657 (no plasmid), 638 (Opto-CheY$_{N449S_{higher}}$), and 658 (CheY-cpAsLOV2$_{N449S}$). Binary traces of cell body rotation (CCW=0, CW=1) are shown as a running average of 100 frames (500 msec). Blue light pulses are shown as vertical blue lines.

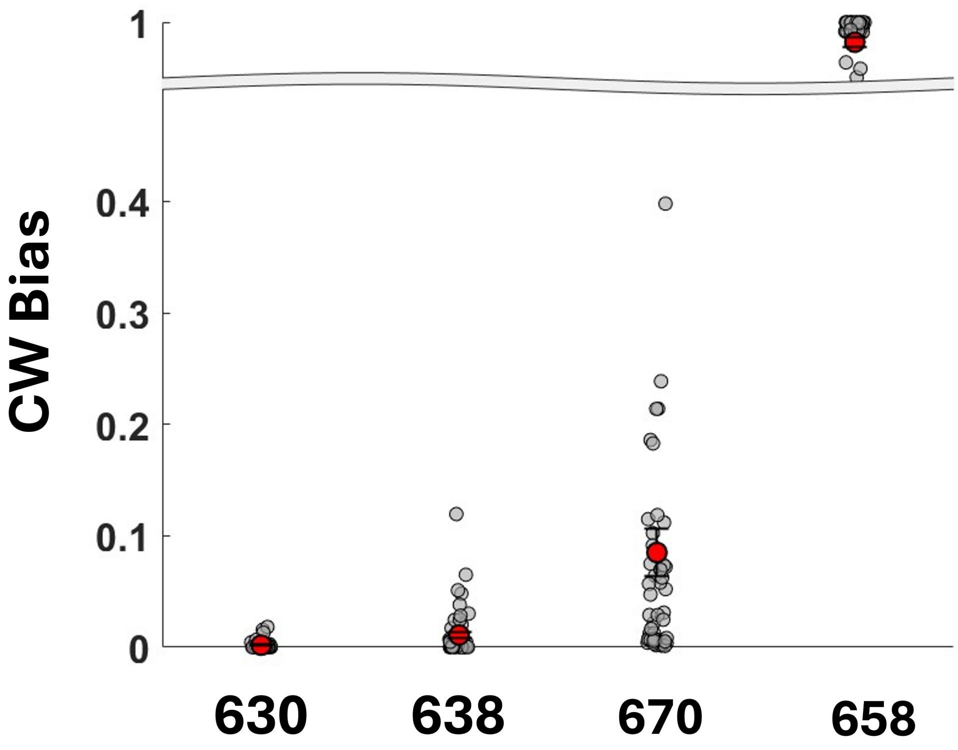


**Fig. S3. Initial CW bias.** Scatter plot of initial CW biases for strains 630, 638, 670, and 658 (Table S1). Note the broken Y-axis. Individual measurements (gray), and means (red) with standard errors are shown. Mean CW biases are 0.00185 $\pm$ 0.00064, 0.011 $\pm$ 0.0032, 0.085 $\pm$ 0.022 and 0.9821 $\pm$ 0.0035, respectively.

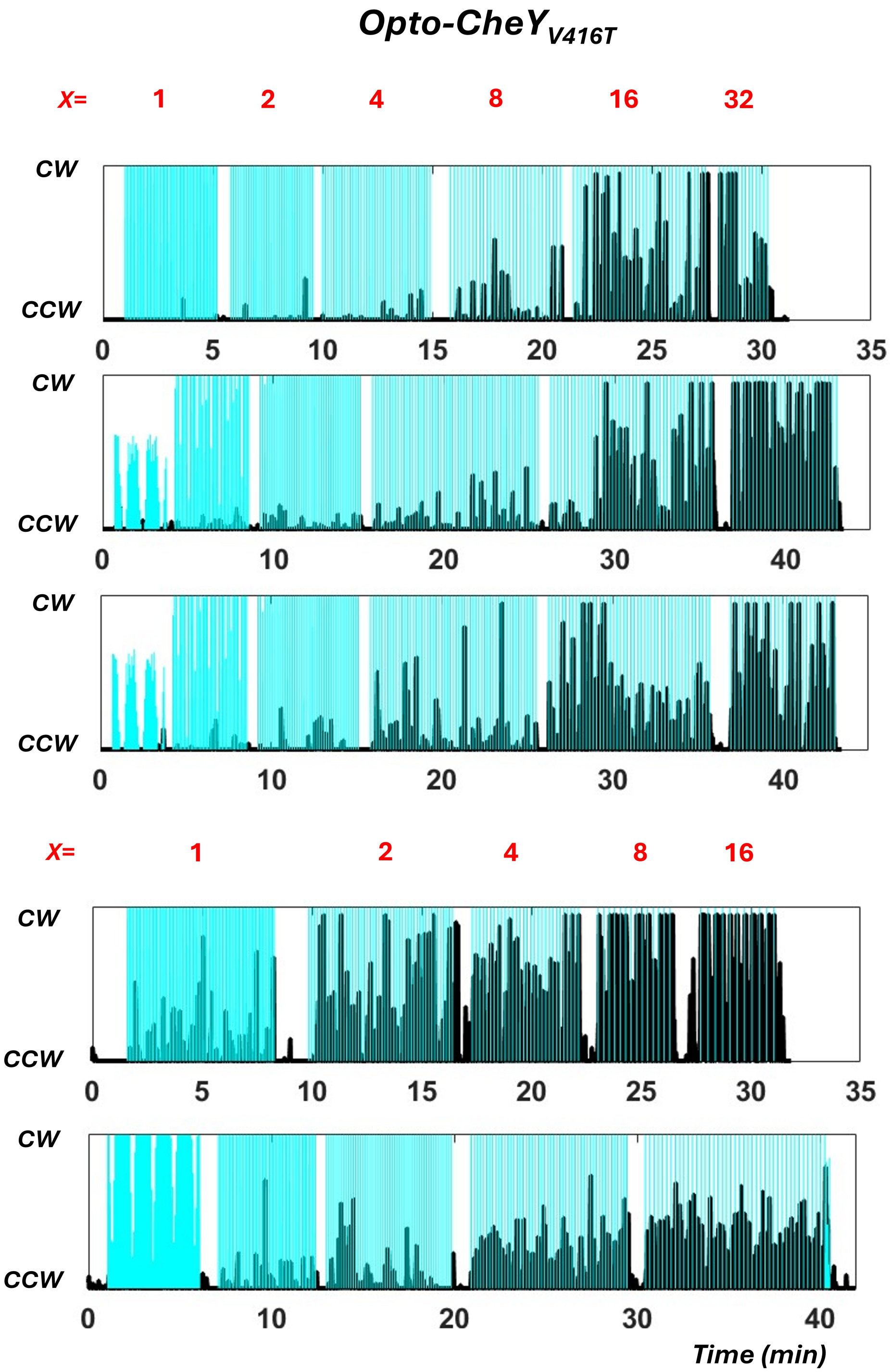


**Fig. S4. Motor responses to light pulses.** Averaged binary traces (100 frames running average, 500 msec) for strain 636 (Opto-CheY$_{V416T}$). Each plot is a single motor trace. Blue light pulses are shown as blue lines. 2 msec blue light pulses of a given blue light intensity ($I = I_{min} \cdot x$, $x = \{1, 2, 4, 8, 16, 32\}$) are applied at constant frequency: every 4 sec for $x = 1$, every 6 sec for $x = 2$, every 8 sec for $x = 4$, every 10 sec for $x = 8$, every 12 sec for $x = 16$, and every 15 sec for $x = 32$.

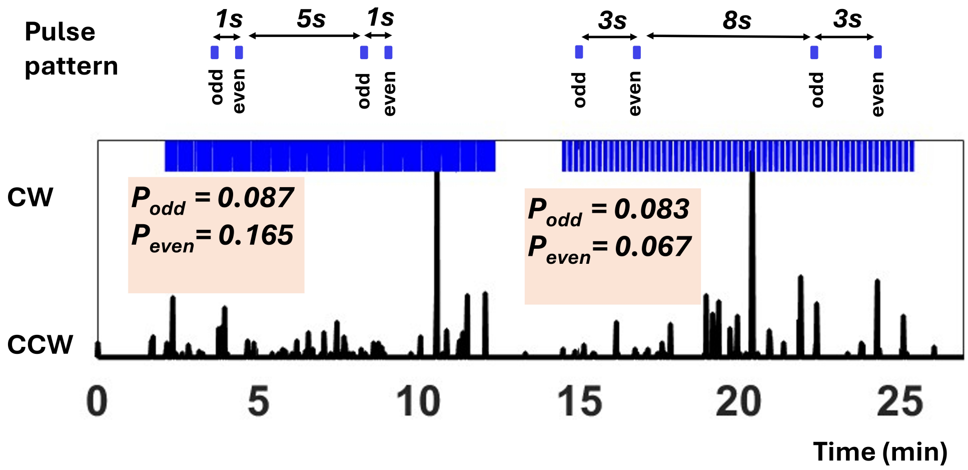


**Fig. S5. Response independence.** Blue light pulses ($x = 4$) are delivered in patterns indicated above the individual motor trace (CW bias averaged with a 500 msec window). 103 doublets of pattern 1, and 60 doublets of pattern 2 were delivered. The probability of motor response was determined for the odd numbered pulses and, respectively, for the even numbered pulses. For pattern 1, $P_{odd}$=0.087, while $P_{even}$=0.165; for pattern 2, $P_{odd}$=0.083 and $P_{even}$=0.067.

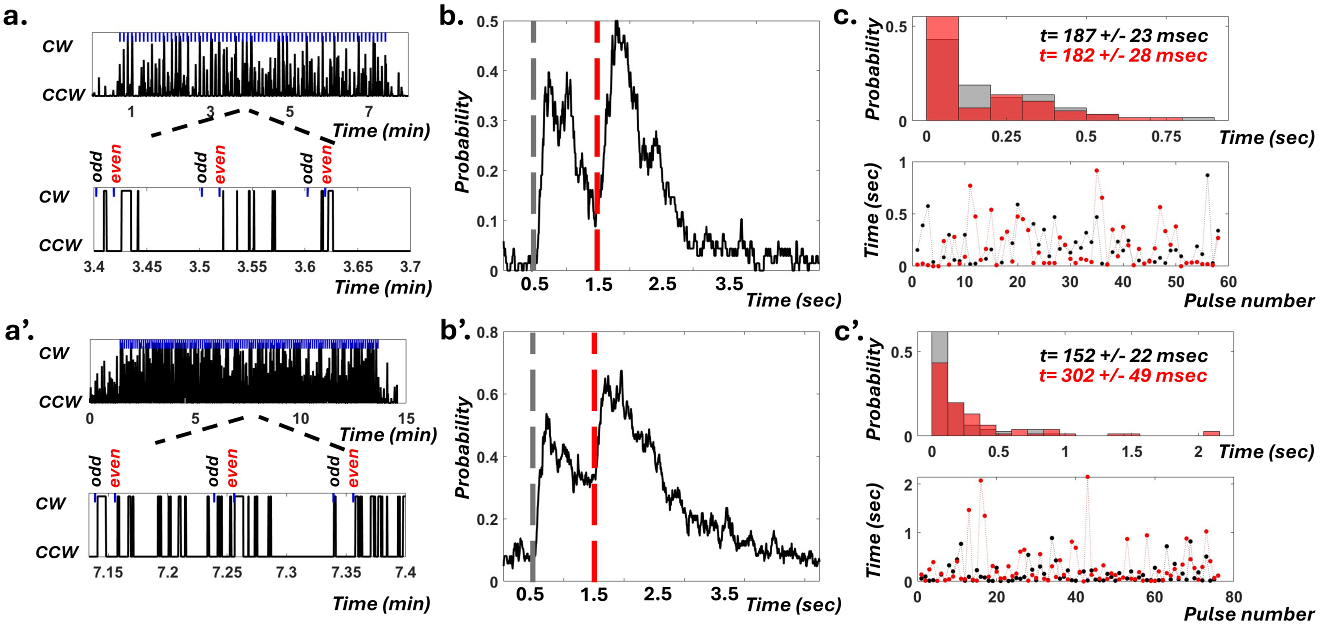


**Fig. S6.** This figure shows data from two different cells (a, b, c), and (a', b', c') with different initial CW average biases (0.03 and 0.06). Panels a/a' show the cell's CW bias as a function of time (100 frames (500 msec) running average for top panel and not averaged for the bottom zoom in). Blue light pulses are marked in blue. Panels b/b' align and average all doublet stimulations giving the probability of motor response for the doublet stimulation as a function of time. The odd pulses are marked in gray, and the even pulses are marked in red. Note that odd pulses happen when the initial CW bias is recovered, while the even pulses happen when the motor's bias is higher than the initial bias. Panels c/c' show histograms for the duration of first CW interval following the odd pulses (gray), and even pulses (red). Average durations are also shown, as well as a plot of the first CW duration as a function of doublet pulse number (successive odd (gray) and even (red) pulses have the same doublet number).

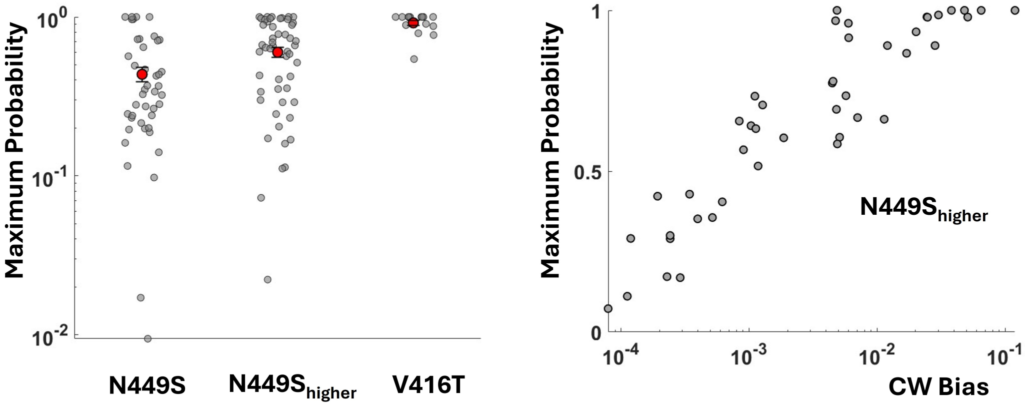


**Fig. S7. Saturating probability of the motor response. Left.** Scatter plot of the maximal probability of motor response for Opto-CheY$_{N449S}$, Opto-CheY$_{N449S_{higher}}$, and Opto-CheY$_{V416T}$ motor (note logarithmic scale). Individual measurements (gray), and means (red) with standard errors are shown. Mean saturating probabilities are 0.435 $\pm$ 0.0435, 0.6 $\pm$ 0.0435 0.0437, and 0.92 $\pm$ 0.034, respectively. **Right.** Maximum response probability as a function of initial CW bias for 51 Opto-CheY$_{N449S_{higher}}$ motors.

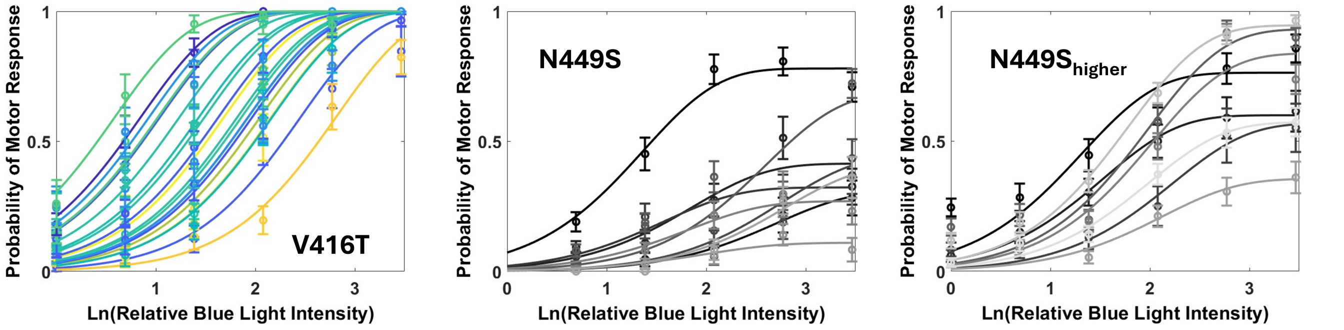


**Fig. S8. Blue light dose response curves for Opto-CheY variants.** Opto-CheY$_{V416T}$, Opto-CheY$_{N449S}$, and Opto-CheY$_{N449S_{higher}}$ single motor dose response curves, as indicated. Each motor responds to light pulses of increasing relative intensity ($I = I_{min} \cdot x$, $x = \{1, 2, 4, 8, 16, 32\}$) controlled using neutral density filters. Response probability for each BFM expressing an Opto-CheY variant at a given blue light intensity is calculated as the number of responses divided by the number of flashes. $P_{see}$ with $\theta = 2$ fits are shown for each of the 20 Opto-CheY$_{V416T}$ dose response measurements in corresponding color. $R^2$ values for the fits are given in Table S5. For Opto-CheY$_{N449S}$ strains, data is fitted to $b\times$ $P_{see}$, where $b$ is between 0 and 1, a free parameter to allow response probability saturation at values less than 1.

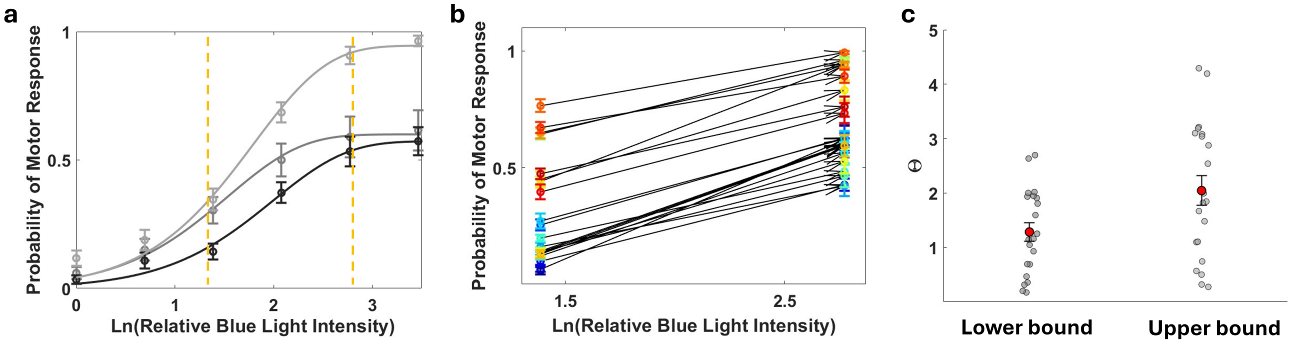


**Fig. S9. Response analysis for Opto-CheY$_{N449S_{higher}}$. a.** The dose response for three individual Opto-CheY$_{N449S_{higher}}$ motors is shown with corresponding $\theta = 2$ fits. Vertical orange lines highlight the intensity interval with most rise in response probability ($x = 4 - 16$). **b.** 25 motors were stimulated at $x = 4$ ($\sim$ 300 flashes) and $x = 16$ ($\sim$ 100 flashes) intensities with enough trials to improve on response probability estimates. For each motor, the response probability estimates are connected by an arrow pointing towards the response probability for $x = 16$. **c.** $\theta$ can be obtained from the slope of the response as $\theta = 2 \cdot \pi \cdot (\frac{\delta P}{\delta ln I})^2$, derivative evaluated at half maximal response. For each motor we can calculate the slope of the line connecting the $x = 4$ and $x = 16$ response probabilities. **c.** The lower bound $\theta$ estimate is made under the assumption that the response is approximately linear in the $x = 4 - 16$ interval, as it is the case for the black colored response in panel **a**. The upper bound estimate is for linear response in the $x = 4 - 8$ interval followed by saturation, as it is the case of the middle gray response in panel **a**. The upper bound estimate is for a linear response in the $x = 4 - 8$ interval followed by saturation, as it is the case of the middle gray response in panel **a**. $\theta$ estimates and corresponding initial CW biases of the motors are anti correlated, the higher the bias, the lower the $\theta$ estimate (Pearson correlation coefficient $r = -0.7622$, and $p = 9.5038e - 06$).

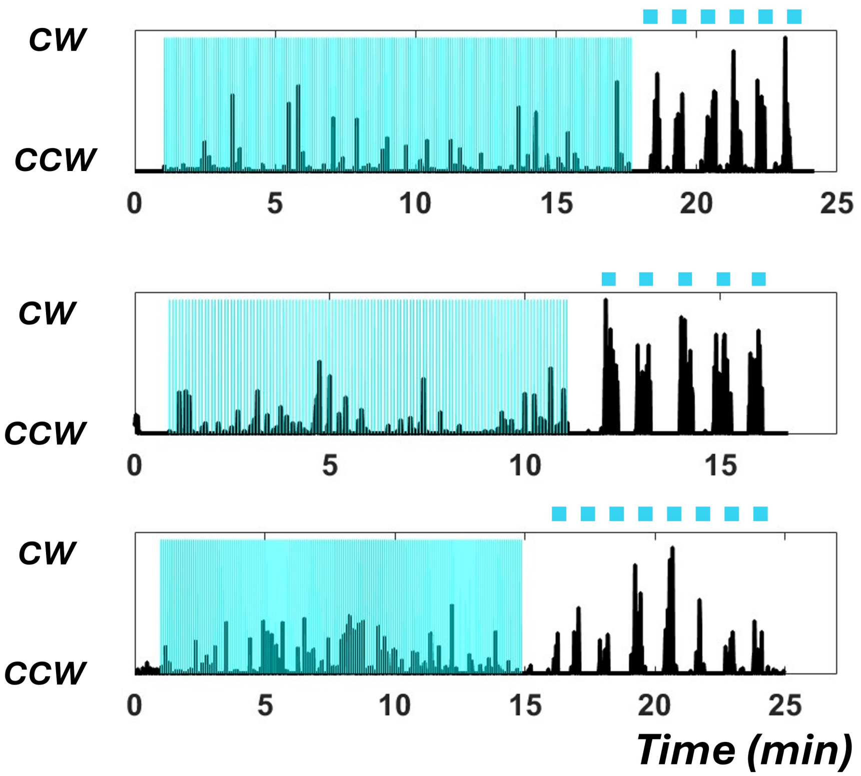


**Fig. S10. Longer pulses increase response probability.** Averaged binary traces (100 frames running average, 500 msec) are shown for strain 630 (Table S1). Tethered cells (Opto-CheY$_{N449S}$) are subjected to 2 msec pulses every 6 sec at the highest intensity (vertical blue lines). Following are 20 sec pulses (horizontal blue lines) at $1/32$ of the maximum intensity.

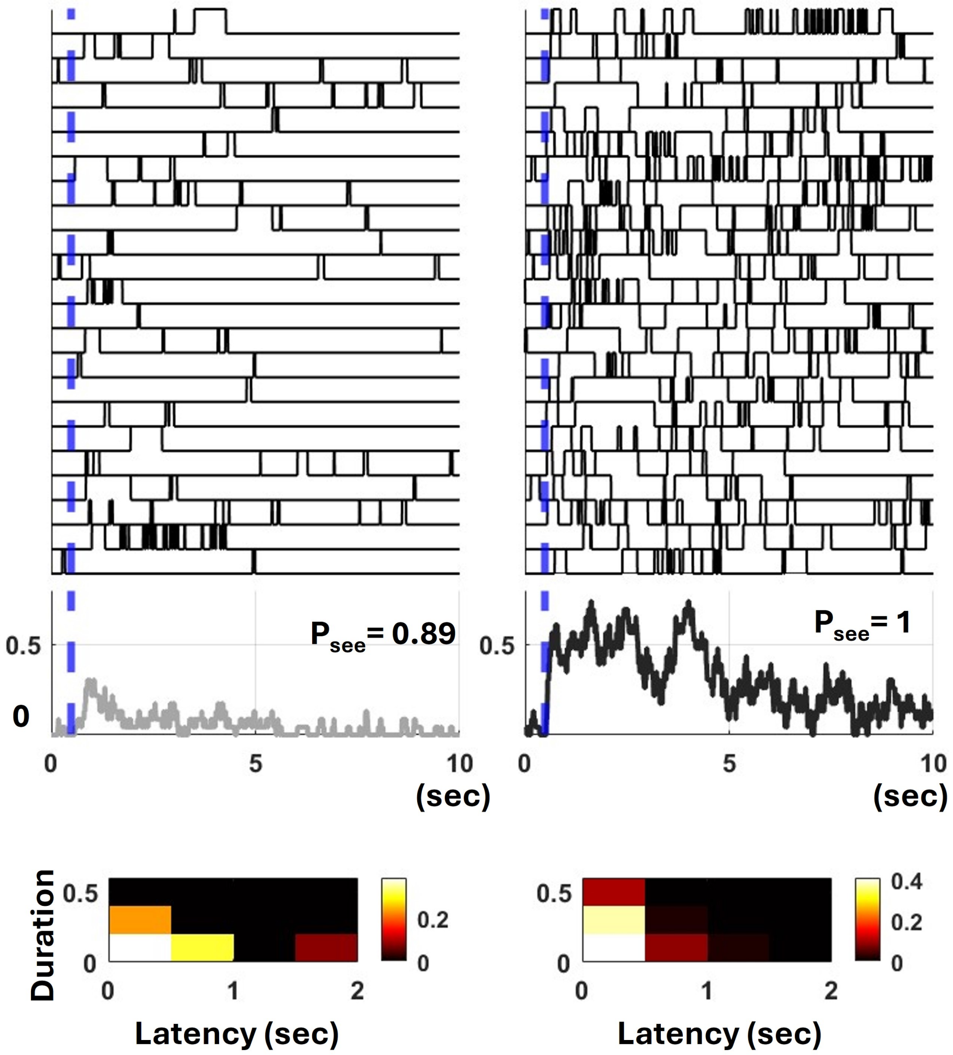


**Fig. S11. The CheY$_{V416T}$ motor responses to different strength stimuli.**. A single motor is subjected to 28 light pulses ($I = I_{min} \cdot x$, $x = 4$, weak stimulus), and 88 pulses ($x = 16$, strong stimulus) 23 responses are stacked and aligned relative to the light pulse (blue line). Average motor responses for each signal strength is shown (gray for the weak stimulus, and black for the strong stimulus). Below, the 23 latencies and durations are used to generate a joint probability distribution as a 2D-histogram, and color coded for probability as shown in the colorbar.

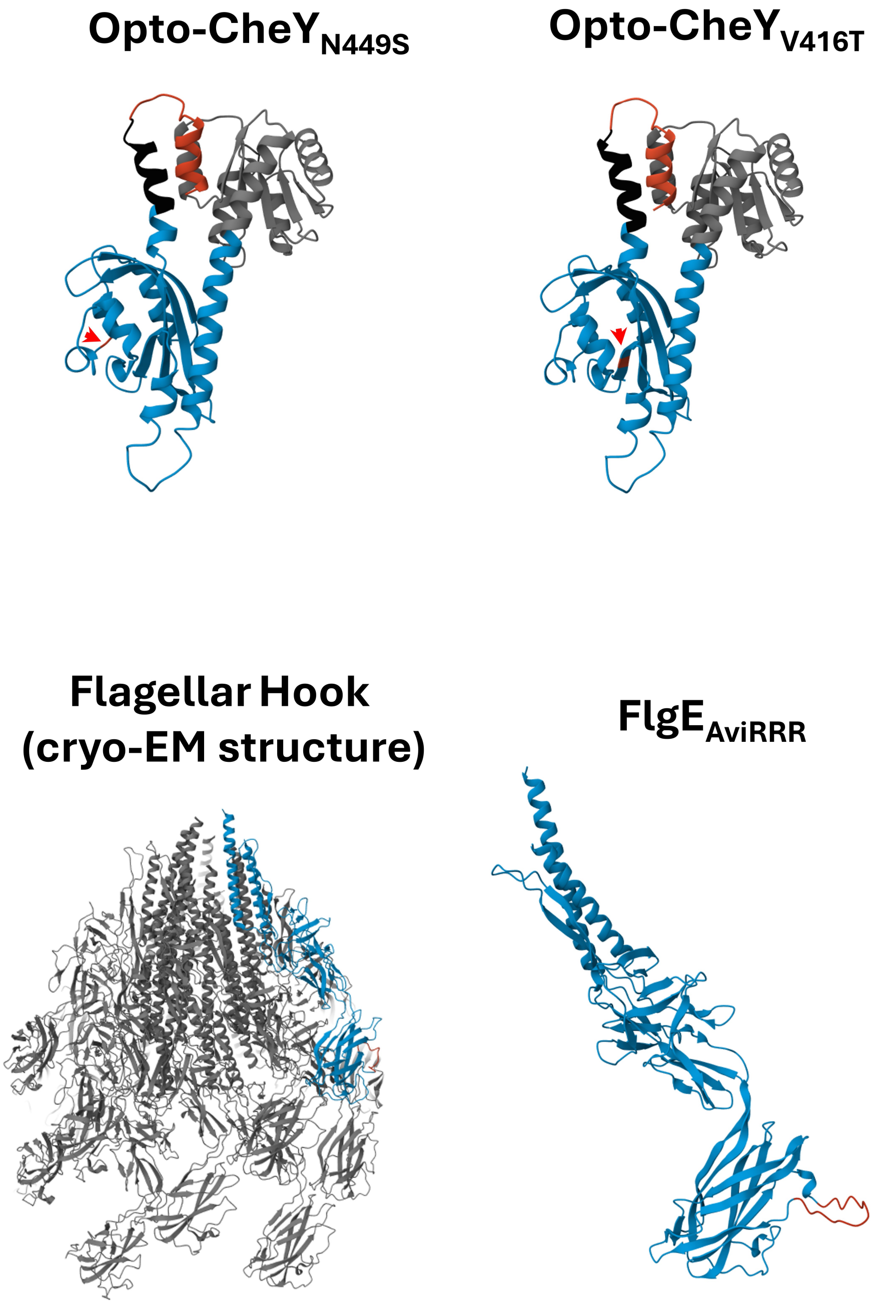


**Fig. S12. Structure predictions for engineered proteins.** AlphaFold first ranked model predictions are shown for Opto-CheY$_{N449S}$ and Opto-CheY$_{V416T}$, color scheme as in Fig. 1b. Red arrow heads point to the location of the introduced mutations. Below, the cryo-EM structure of the flagellar hook from *Salmonella* is shown (pdb ID 7CGB).(4) One FlgE subunit of the hook is highlighted in blue. AlphaFold structure prediction of FlgE$_{AviRRR}$ is shown to the right (engineered loop in red).

### Movies

Tethered cells movies are shown for strains 630, 636, 638, 657, 658 (Table S1). Each video shows 5 responses for the same cell, vertically stacked and aligned relative to the blue light pulse (2 msec, frame 101). For strain 636 , blue light intensity is at 1/2 of the maximal intensity (~50 $\mu$J/cm$^2$, every 12 seconds (2400 frames)), while for the rest blue light intensity is maximal (~100 $\mu$J/cm$^2$, every 6 sec (1200 frames)). Videos are acquired at 200 fps, but shown here at 40 fps. A synchronized plot that tracks the rotational direction of the cell as a function of frame number is shown next to each video.

Movie 1. A 657 tethered cell.
Movie 2. A 638 tethered cell.
Movie 3. Another 638 tethered cell.
Movie 4. A 658 tethered cell.
Movie 5. A 630 tethered cell.
Movie 6. Another 630 tethered cell.
Movie 7. 636 tethered cell.
Movie 8. Another 636 tethered cell.
